# STI1 domain dynamically engages transient helices in disordered regions to drive self-association and phase separation of yeast ubiquilin Dsk2

**DOI:** 10.1101/2025.03.14.643327

**Authors:** Nirbhik Acharya, Emily A. Daniel, Thuy P. Dao, Jessica K. Niblo, Erin Mulvey, Shahar Sukenik, Daniel A. Kraut, Jeroen Roelofs, Carlos A. Castañeda

**Affiliations:** Department of Biology, Syracuse University, Syracuse, NY 13244, USA; Department of Chemistry, Syracuse University, Syracuse, NY 13244, USA; Department of Biochemistry and Molecular Biology, University of Kansas Medical Center, Kansas City, KS 66160, USA; Department of Chemistry, Villanova University, Villanova, PA 19085, USA; BioInspired Institute, Syracuse University, Syracuse, NY 13244, USA; Interdisciplinary Neuroscience Program, Syracuse University, Syracuse, NY 13244, USA

**Keywords:** transient helices, intrinsically disordered regions, STI1 domain, phase separation, proteasome condensates, yeast, ubiquilins

## Abstract

Ubiquitin-binding shuttle proteins are important components of stress-induced biomolecular condensates in cells. Yeast Dsk2 scaffolds proteasome-containing condensates via multivalent interactions with proteasomes and ubiquitinated substrates under azide-induced mitochondrial stress or extended growth conditions. However, the molecular mechanisms underlying how these shuttle proteins work are unknown. Here, we identify that the middle chaperone-binding STI1 domain is the main driver of Dsk2 self-association and phase separation *in vitro*. Using NMR spectroscopy and computational simulations, we find that the STI1 domain interacts with three transient amphipathic helices within the intrinsically-disordered regions of Dsk2. Removal of either the STI1 domain or these helices significantly reduces the propensity for Dsk2 to phase separate. *In vivo*, removal of the STI1 domain in Dsk2 has the opposite effect, resulting in an increase of proteasome-containing condensates due to an accumulation of polyubiquitinated substrates. Modeling of STI1-helix interactions reveals a binding mode that is reminiscent of interactions between chaperone STI1/DP2 domains and client proteins containing amphipathic or transmembrane helices. Our findings support a model whereby STI1-helix interactions important for Dsk2 condensate formation can be replaced by STI1-client interactions for downstream chaperone or other protein quality control outcomes.

**Highlights:**

- The intrinsically disordered regions of Dsk2 harbor transient helices that regulate protein properties via interactions with the STI1 domain.
- The STI1 domain is a significant driver of Dsk2 self-association and phase separation *in vitro*.
- Dsk2 colocalizes with ubiquitinated substrates and proteasome in reconstituted condensates.
- Absence of Dsk2 STI1 domain in stressed yeast cells promotes formation of proteasome condensates coupled with upregulation of polyubiquitinated substrates.

## Introduction

Ubiquitin (Ub)-binding shuttle proteins and adaptors are important drivers and components of biomolecular condensates for protein quality control, stress response, and immune system activation (Yasuda *et al*, 2020; Uriarte *et al*, 2021; Waite *et al*, 2024; Zaffagnini *et al*, 2018; Sun *et al*, 2018; Rajendran & Castañeda, 2025; Goel *et al*, 2023; Du *et al*, 2022). A subset of these Ub-binding shuttle proteins, called ubiquilins (UBQLNs), form condensates with ubiquitinated substrates, proteasomes, and/or other PQC machinery under either physiological or stress-induced conditions (Dao *et al*, 2018; Alexander *et al*, 2018; Waite *et al*, 2024; Gerson *et al*, 2021; Mohan *et al*, 2022, 2024). Dysregulation of human UBQLNs is linked to neurodegeneration (Deng *et al*, 2011; Lin *et al*, 2022; Edens *et al*, 2017). In other organisms, including yeast and plants, removal of UBQLN orthologs negatively affects organismal stress responses (Waite *et al*, 2024; Nolan *et al*, 2017; Chuang *et al*, 2016; Jantrapirom *et al*, 2020). In yeast, Dsk2, the ortholog of human UBQLNs, and Rad23 are the two main shuttle factors of the ubiquitin proteasome system (UPS) (Samant *et al*, 2018; Tsuchiya *et al*, 2017). Dsk2 and Rad23 are critical drivers of proteasome condensates in stressed yeast cells and deletion of the genes encoding both proteins abrogates the formation of proteasome-containing condensates in yeast (Waite *et al*, 2024). However, the molecular underpinnings of this process are unknown.

Dsk2 (373 amino acids (AA)) shares a similar domain architecture with the human UBQLNs (Figure S1). Each contains UBL (Ub-like) and UBA (Ub-associating) domains that interact with proteasome receptors and ubiquitinated substrates, respectively (Figure 1A) (Funakoshi *et al*, 2002). In-between the folded UBL and UBA domains are predicted intrinsically-disordered regions (IDRs) and a single STI1-like domain (as opposed to two STI1 domains in UBQLNs). STI1 domains engage with chaperones and mediate interactions with client proteins, such as transmembrane domains of mitochondrial proteins (Itakura *et al*, 2016; Schmid *et al*, 2012; Lin *et al*, 2021; Onwunma *et al*, 2024). However, there is limited structural and dynamical information about STI1 domains and their binding mechanisms (Fry *et al*, 2021). AlphaFold predictions of STI1 domains are generally of low confidence and show non-compact structures. Recent work from our lab demonstrated that the STI1-II domain of UBQLN2 is important for mediating both self-association and phase separation (Dao *et al*, 2018, 2024). Several ALS/FTD disease-linked mutations in UBQLN2 localize to the STI1 domains, but little is known about their underlying disease mechanisms (Renaud *et al*, 2019).

**Figure 1.**
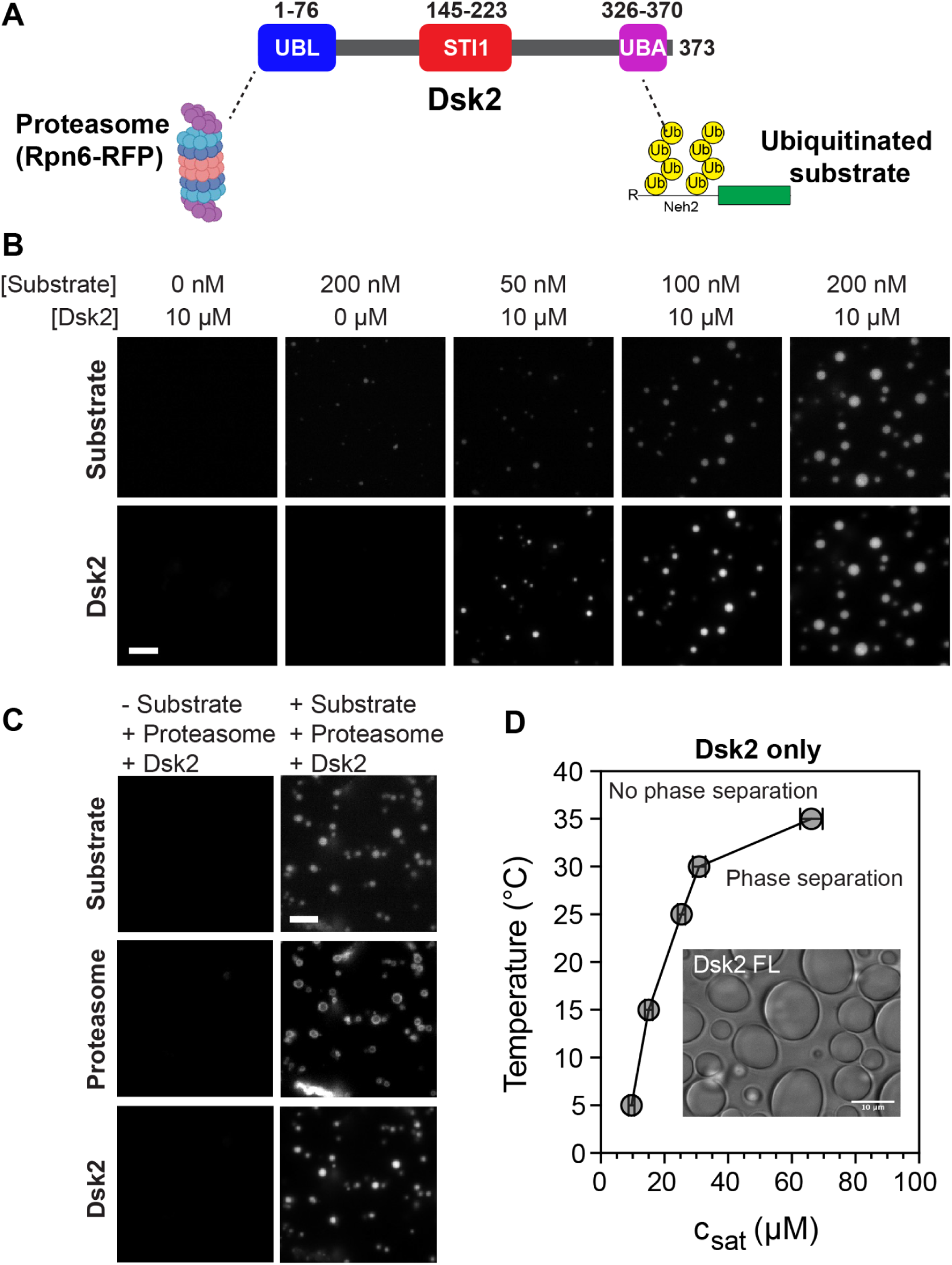
Dsk2 forms condensates with proteasomes and ubiquitinated substrates. (A) Domain architecture of Dsk2 highlighting known domain interactions with proteasomes and ubiquitinated substrates (Ub-substrate). (B) Fluorescence microscopy showing phase separation of Ub-substrate with Dsk2 in a concentration-dependent manner (3% PEG8K, 18°C, 0 or 10 µM Dsk2, 0-200 nM substrate). Scale bar, 5 µm. (C) Fluorescence microscopy of 10 µM Dsk2, 200 nM Ub-substrate (absence or presence), and 100 nM proteasome incubated at 18°C in pH 7.5 buffer with 3% PEG8K. Scale bar, 5 µm. (D) Temperature-concentration phase diagram of Dsk2 (alone) delineating the saturation concentration (c_sat_) at different temperatures in buffer containing 20 mM NaPhosphate pH 6.8, 150 mM NaCl, 7.5% PEG8K, 0.5 mM EDTA. Inset: brightfield microscopy of 100 µM Dsk2 droplets incubated at 25°C in 20 mM NaPhosphate pH 6.8, 150 mM NaCl, 7.5% PEG8K, 0.5 mM EDTA. Error bars represent the standard deviation over three protein preps performed in triplicates.

Here, we demonstrate that the STI1 domain is a major contributor for the self-association and phase separation of Dsk2 *in vitro*, in a similar role as the STI1-II domain to UBQLN2. Critically, we obtain residue-by-residue information using NMR (nuclear magnetic resonance) spectroscopy for >84% of full-length Dsk2. We map interactions across Dsk2 and determine that the STI1 domain engages with transient helices within disordered regions of Dsk2. We reconstitute condensates containing Dsk2, ubiquitinated substrates, and proteasomes that complement previous results in stressed yeast cells. We demonstrate that in stressed yeast cells the deletion of the STI1 domain causes accumulation of polyubiquitinated proteins, confounding the impact on proteasome condensates. Our data suggest that the interaction between the STI1 domain and transient helices in Dsk2 may represent a conserved feature among related UBQLNs and co-chaperone like proteins containing STI1-like domains (Sti1, HIP, SGTA, Tic40).

## Results

### Dsk2 phase separates under physiological conditions

Ub-binding shuttle proteins Dsk2 and Rad23 are required for proteasome condensate formation under stress conditions in yeast (Waite *et al*, 2024). To determine the role of Dsk2 in proteasome condensates, we reconstituted a minimal system *in vitro* using only Dsk2, K63-linked ubiquitinated substrate (Ub-substrate), and yeast proteasome (Figure 1A). Using fluorescence microscopy, we tested and determined that Ub-substrate phase separates with Dsk2 in a concentration-dependent manner, consistent with our recent results on UBQLN2 and ubiquitinated substrate (Figure 1B) (Valentino *et al*, 2024). Furthermore, we also observed colocalization of proteasome, Dsk2, and Ub-substrate in droplets (Figure 1C), consistent with previous observations in yeast (Waite *et al*, 2024). In the absence of the proteasome and Ub-substrate, we found that Dsk2 phase separates in the presence of a higher amount of macromolecular crowder polyethylene glycol (PEG) at sufficiently high protein concentrations (Figure 1D). We obtained a phase diagram of Dsk2 by measuring the saturation concentration (c_sat_) of Dsk2 as a function of temperature (see Methods). Upon decreasing temperature, Dsk2 phase separates at concentrations reaching 10 µM at 5°C; these data are consistent with Dsk2 following an upper critical solution temperature (UCST) phase transition whereby decreasing temperature favors phase separation.

### Short helices exist within intrinsically disordered regions of Dsk2

To map out different structural elements within Dsk2, we used biomolecular NMR spectroscopy as previously employed in (Dao *et al*, 2018, 2022; Conicella *et al*, 2016). We expressed and purified ^15^N full-length Dsk2 (Dsk2 FL) protein (Methods, Figure S2). We collected ^1^H-^15^N HSQC NMR spectra (Figure 2A) to monitor backbone amide resonances on a residue-by-residue level at 50 µM protein concentration, where Dsk2 is a monomer according to size exclusion chromatography - multi-angle light scattering (SEC-MALS) experiments (Figure S3). A majority of amide resonances are in the middle of the spectrum (7.5-8.5 ppm in ^1^H), consistent with the presence of intrinsically disordered regions (IDRs) in Dsk2 (Dao *et al*, 2018; Burke *et al*, 2015). Peaks located on either side of this middle region correspond to residues in the well-folded UBL and UBA domains, as previously determined (Zhang *et al*, 2009). Using triple resonance NMR experiments with several domain deletion variants of Dsk2, we obtained chemical shift assignments for 84% of all Dsk2 FL backbone amide resonances (see Methods/SI). Most of the missing peaks correspond to resonances in the STI1 domain, suggesting increased dynamics within the STI1 domain. We probed secondary structure content using ^13^C backbone C_α_ chemical shifts (Figure 2B). Our experimentally determined C_α_ values for Dsk2 correlate very well with predicted C_α_ values determined from machine learning and the Alphafold-derived structure of Dsk2 (Figure 2B, 2C; see Methods) (Gu *et al*, 2024). The STI1 domain is largely helical with the magnitude of C_α_ secondary shifts comparable to C_α_ values for the well-structured helices in the UBA domain. By contrast, the regions in between the UBL, STI1, and UBA domains are largely disordered. The ΔδC_α_ plot also captures three regions (114-134, 279-291 and 303-313) in IDRs with helical propensity, indicating presence of putative α-helices.

**Figure 2.**
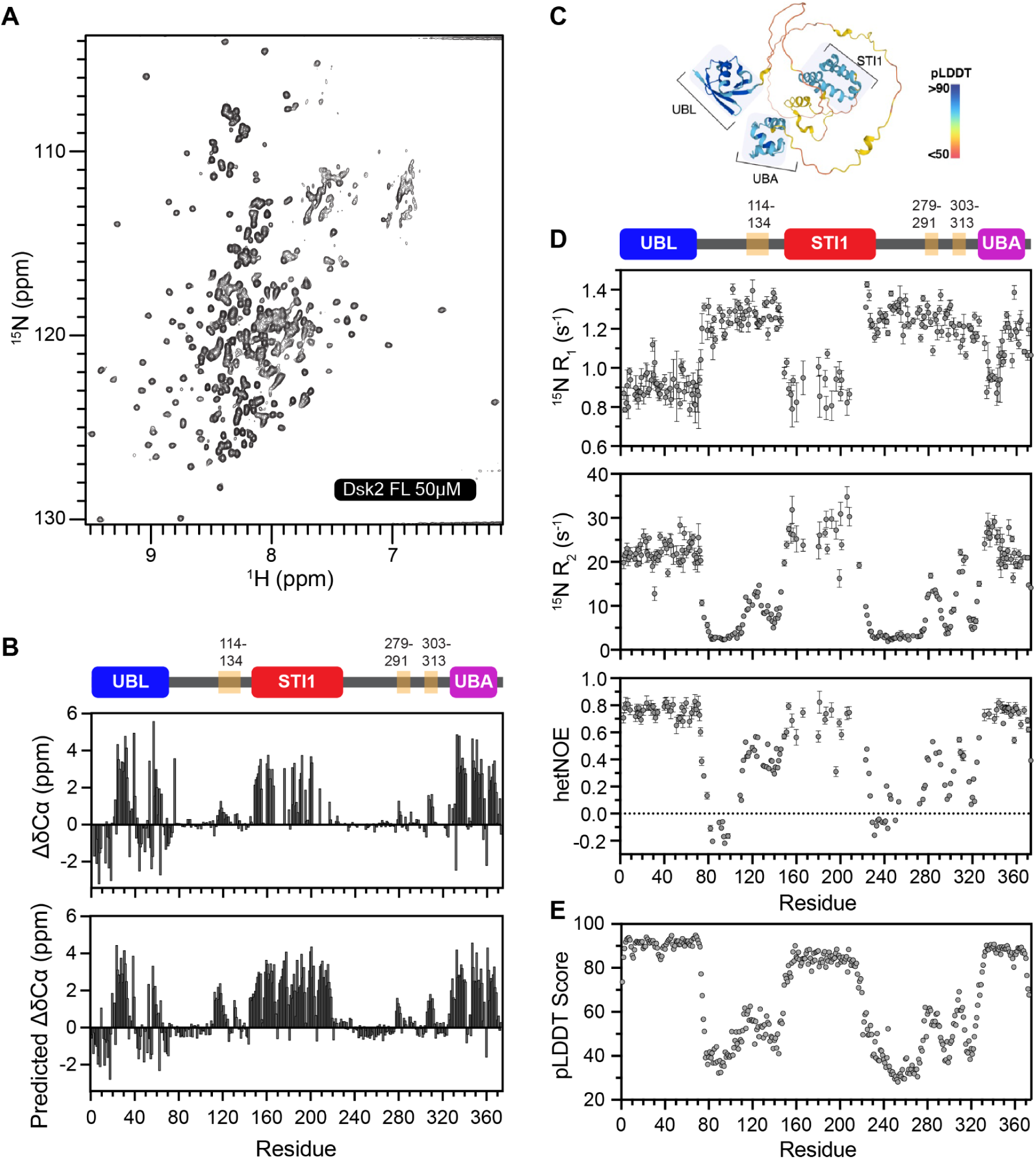
Dsk2 is a multidomain protein with three transient helical regions in intrinsically disordered regions. (A) ^1^H-^15^N TROSY-HSQC spectra of 50 µM Dsk2 FL at 25°C in NMR buffer (20 mM NaPhosphate, 0.5 mM EDTA, 0.02% NaN3 pH 6.8). (B) Residue-level secondary structure comparison between experimentally determined C_α_ chemical shifts and EFG-CS predicted C_α_ chemical shifts from Dsk2 AlphaFold structure (See Methods). Positive and negative ΔδC_α_ values indicate α-helix and β-sheet content, respectively. (C) AlphaFold-predicted structure of Dsk2. (D) Amide backbone ^15^N R_1_ relaxation rates, ^15^N R_2_ relaxation rates, and {^1^H}-^15^N hetNOE values for Dsk2 FL at 50 µM. Errors in R_1_ and R_2_ were determined using 500 Monte Carlo trials using RELAXFIT on MATLAB. Errors in hetNOE measurements were determined using the SE propagation formula. (E) AlphaFold-predicted local distance difference test (pLDDT) score for Dsk2.

To map backbone amide dynamics on a residue-by-residue level, we collected standard NMR ^15^N longitudinal (R_1_) relaxation, ^15^N transverse (R_2_) relaxation, and {^1^H-^15^N} heteronuclear Overhauser enhancement (hetNOE) experiments (Figure 2D). R_1_ relaxation rates monitor nanosecond-picosecond backbone dynamics, while R_2_ relaxation rates are also sensitive to slower millisecond-microsecond backbone dynamics. hetNOE reports on local dynamics at the picosecond-nanosecond scale. Resonances within the UBL, STI1, and UBA domains exhibited high hetNOE values (>0.6), consistent with these residues being part of structured regions. Elevated R_2_ rates and lowered R_1_ rates among these residues suggest similar rotational tumbling times, likely due to these domains being of similar sizes. In contrast, resonances within disordered regions have low R_2_ relaxation rates (2-4 s^-1^) and low hetNOE values (∼0.1 or lower) similar to previously reported data for IDRs (Dao *et al*, 2018; Burke *et al*, 2015). Notably, the three putative helical regions have intermediate hetNOE values (∼0.2-0.4) suggestive of their transient helical character (hereafter referred to as transient helices (TH)). Remarkably, the pattern of R_2_ relaxation rates and hetNOE values closely matched the AlphaFold pLDDT scores across the entire protein (Figure 2E). High pLDDT scores indicate strong confidence in the structural prediction, and these regions correspond to well-folded domains in Dsk2. All three transient helices (hereafter 3TH when referring to all three regions together) exhibit intermediate pLDDT scores, consistent with their intermediate hetNOE values and R_2_ relaxation rates. Together, these data show that the UBL, STI1, and UBA domains are well-folded, with the 3TH regions exhibiting transient helical propensity.

### Multivalent interactions drive Dsk2 self-association

Macromolecular phase separation can be driven by multivalent interactions among folded and/or disordered regions in proteins (Pappu *et al*, 2023; Martin *et al*, 2021). Using SEC-MALS, we determined that Dsk2 predominantly exists as a monomer at 150 µM, but we do observe a decrease in the elution volume as protein concentration is increased to 300 µM. This indicates that Dsk2 forms dynamic self-associating assemblies as protein concentration increases (Figure S3). To assess the residue-level contribution of different domains/regions in Dsk2 self-association, we measured concentration-dependent effects by comparing ^1^H-^15^N TROSY-HSQC NMR spectra of FL Dsk2 between 50 µM and 400 µM, representing monomer and dynamic self-association, respectively (Figure 3A). Changes to NMR spectra at 400 µM stem from both inter- and intra-molecular interactions. Strikingly, at high protein concentration, many peaks corresponding to residues in the STI1 domain either disappeared or exhibited ∼90% decrease in peak intensity (Figure 3A inset). The peak intensities in the UBL and UBA domains decreased by 50-70%. Intensity changes in UBL and UBA resonances stem from previously-known UBL-UBA interactions common to Ub-binding shuttle proteins (Lowe *et al*, 2006; Zientara-Rytter & Subramani, 2019; Zheng *et al*, 2021). We validated these UBL-UBA interactions by comparing chemical shifts of the UBL or UBA domains in the absence and presence of the rest of the Dsk2 protein (Figure S4). Additionally, there was a moderate decrease in peak intensities and subtle chemical shift perturbations (CSPs) for the 3TH regions (Figure 3B and 3C). The concentration-dependent changes in ^15^N R_2_ relaxation rates also point towards self-association governed by different regions of Dsk2 (Figure S5). Together, these data implicate the UBL, UBA, STI1, and the 3TH regions in Dsk2 self-association.

**Figure 3.**
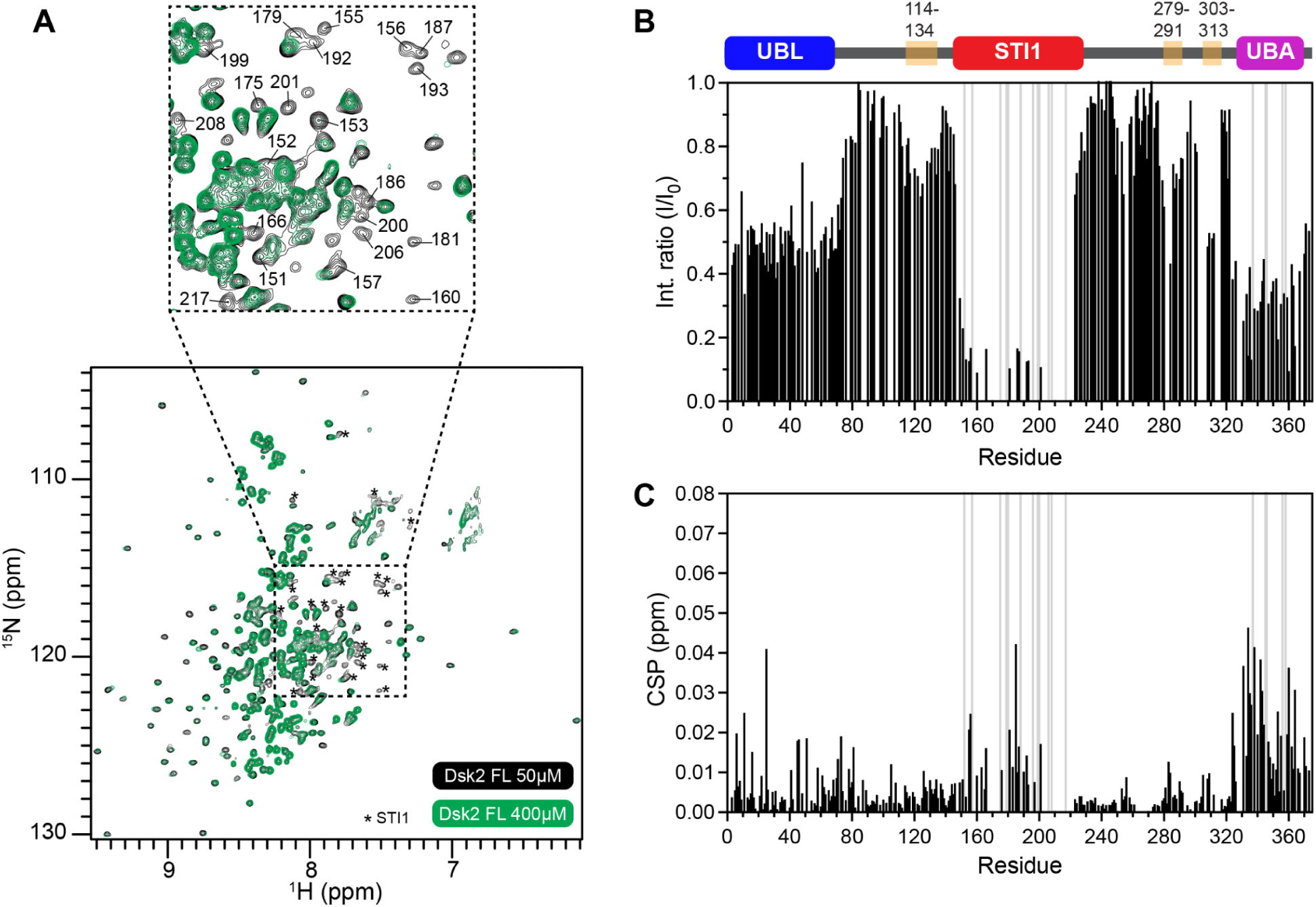
NMR reveals multivalent interactions in Dsk2 self-association. (A) Comparison of ^1^H-^15^N TROSY-HSQC spectra of 50 µM and 400 µM full-length (FL) Dsk2. Inset shows significant line broadening for select amide resonances at 400 µM (green), primarily in the STI1 domain. (B) Residue-specific intensity ratio (I/I_0_) between 50 µM (I) and 400 µM (I_0_) resonances. (C) CSPs represent residue-specific chemical shift differences between 50 µM and 400 µM Dsk2. Gray bars in B and C panels represent resonances with no observable peak at 400 µM protein concentration.

By far, the largest changes in peak intensity occur in STI1 resonances, suggestive of intermolecular STI1-STI1 interactions and/or STI1 interactions with other regions of Dsk2 at an intramolecular or intermolecular level. To test this, we monitored concentration-dependent changes in peak intensity for several domain deletion variants of Dsk2, including ΔUBL and two constructs of Dsk2 lacking both UBL and UBA domains (Figure S6). Resonances corresponding to residues from the STI1 domain consistently exhibited the largest decreases in peak intensity (70-90%) at high protein concentration, suggestive of STI1-STI1 self-association even in the absence of UBL and UBA domains. Our results suggest that the STI1 domain is a major contributor to Dsk2 self-association.

### STI1 domain interacts with transient helices in Dsk2

To investigate how the STI1 domain interacts with the rest of the Dsk2 protein, we made a Dsk2 ΔSTI1 (STI1 deletion) construct and compared ^1^H-^15^N HSQC NMR spectra of Dsk2 FL and Dsk2 ΔSTI1 at identical protein concentrations (50 µM) (Figure 4A). We observed that amide resonances in only the 3TH regions showed large changes in chemical shifts upon STI1 deletion. This impact is quantified in the CSP plot where the 3TH regions show significantly higher CSPs (>0.05 ppm) compared to the rest of the protein (Figure 4B). To fully understand the impact of STI1 domain deletion on 3TH, we compared the backbone dynamics using ^15^N R_1_, R_2_ relaxation rates, and {^1^H-^15^N} hetNOE values (Figure 4C). Strikingly, deletion of the STI1 domain resulted in significant decrease of R_2_ relaxation rates for all three helical regions. The STI1 deletion-induced changes in 3TH backbone dynamics could result from: a) removal of STI1-3TH interactions, or b) destabilization of the secondary structure of these transient helices. However, the latter is unlikely as the measured hetNOE values (Figure 4C) and α-helix propensity (Figure 4D) for the 3TH regions in ΔSTI1 remain similar when compared with data for Dsk2 FL.

**Figure 4.**
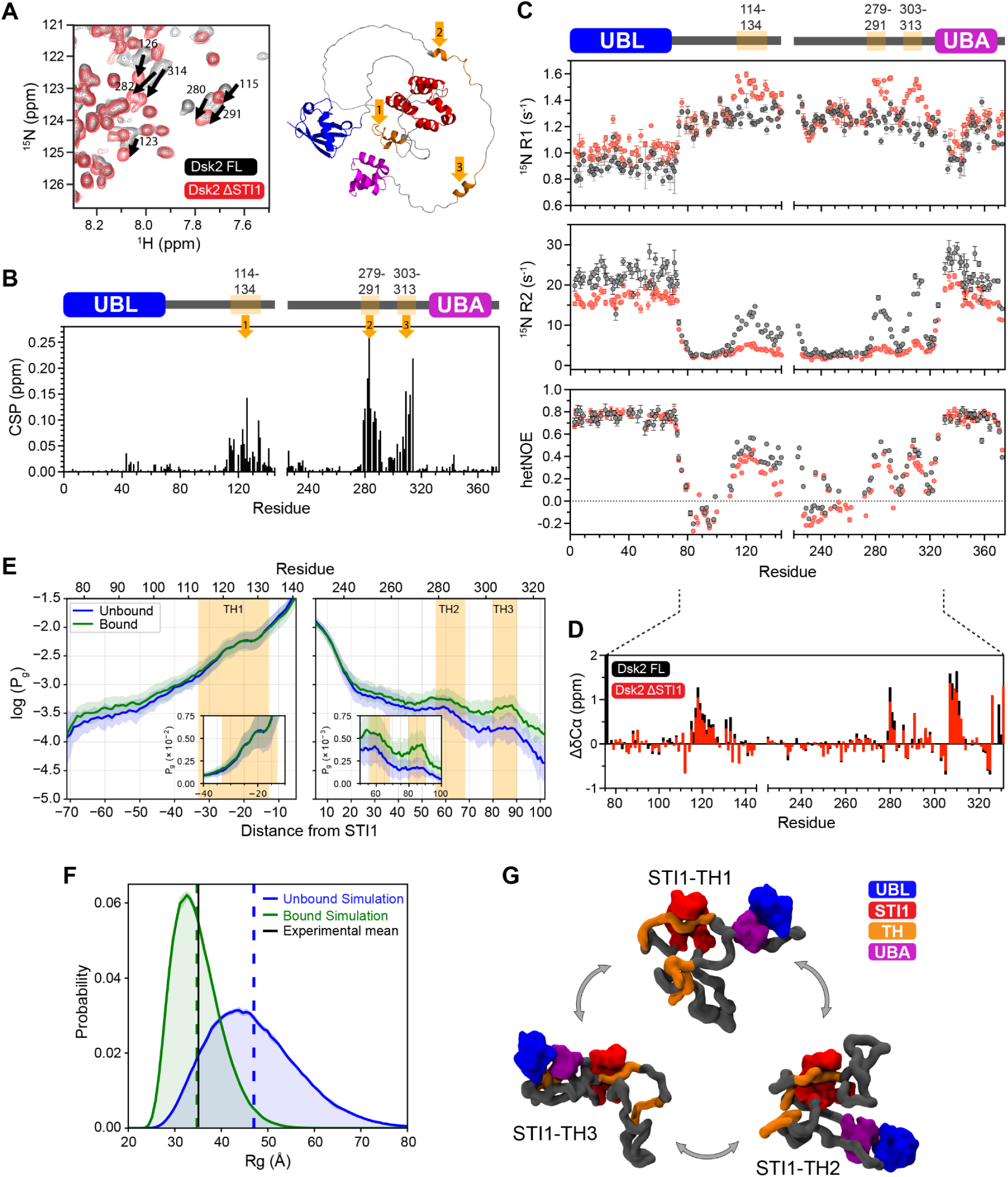
STI1 domain interacts with transient helices present in the IDRs of Dsk2. (A) Comparison of ^1^H-^15^N TROSY-HSQC spectra of 50 µM Dsk2 FL (black) and Dsk2 ΔSTI1 (red) at 25°C focused on amide resonances from the three transient helical regions (3TH) in Dsk2 ΔSTI1; these 3TH are highlighted on the AlphaFold Dsk2 structure on right. (B) Residue-level amide CSPs between Dsk2 FL and Dsk2 ΔSTI1 with largest chemical shift differences for residues from 3TH regions in the IDRs of Dsk2 highlighted by orange arrows and mapped onto Dsk2 AlphaFold structure. (C) ^15^N R_1_, ^15^N R_2_ relaxation rates, and hetNOE values are compared for Dsk2 FL (black) and Dsk2 ΔSTI1 (red) at 50 µM 25°C. Errors in R_1_ and R_2_ were determined using 500 Monte Carlo trials using RELAXFIT on MATLAB. Errors in hetNOE measurements were determined using the SE propagation formula. (D) Residue-specific experimentally determined C_α_ chemical shifts of IDRs are compared between Dsk2 FL and Dsk2 ΔSTI1. (E) Log of the probability of the disordered residues occupying the STI1 groove for CALVADOS simulations where the UBL and UBA were unbound (blue) or bound (green). TH regions marked with orange background. Inset shows the linear probability for the three transient helical regions. The shaded region represents the differences among the replicates. (F) Simulation-derived R_g_ probability distribution compared between the UBL:UBA unbound (blue) and bound (green) forms of Dsk2 with average R_g_ values represented as dashed lines. Experimental R_g_ from SAXS is shown as black solid line. (G) Snapshots from single-protein simulations with the UBL and UBA in a bound conformation showing TH1, TH2, or TH3 occupying the STI1 groove.

To further probe the nature of STI1-TH interactions, we conducted molecular dynamics simulations for full-length Dsk2 using CALVADOS3, a coarse-grained model for disordered and multidomain proteins where each residue is represented as a single bead (Tesei *et al*, 2021; Tesei & Lindorff-Larsen, 2023; Cao *et al*, 2024). In these simulations, the UBL, STI1, and UBA were treated as folded domains held together by elastic constraints (Cao *et al*, 2024), with the rest of the protein being disordered. From these single-protein simulations, we quantified the probability of the disordered regions occupying the STI1 groove. We observed an increased preference of all three TH regions to interact with STI1 (Figure 4E, Movie S1). However, in validating these simulations, we found that the average predicted radius of gyration (R_g_) value for these simulations (47.0 ± 0.2 Å) was much larger than the experimental R_g_ value (35.1 ± 0.2 Å) obtained from small angle X-ray scattering (SAXS) data for full-length Dsk2 (Figure 4F, Figure S3). We hypothesized that this difference stemmed from known UBL-UBA interactions (Lowe *et al*, 2006) that were also observed in our NMR data (see above) but not maintained in these simulations. To test this hypothesis, we placed elastic constraints that held together the UBL and UBA domains in the same orientation as the previously solved crystal structure 2BWE of the two isolated domains (see Methods, Figure S4C). Simulations of the bound model showed quantitative agreement with experimental SAXS data for full-length Dsk2 (R_g_ = 34.8 ± 0.1 Å, Figure 4F, Movie S2). In addition, keeping the UBL and UBA in a bound conformation resulted in a 2-fold increased preference for TH2 and TH3 regions to interact with the STI1 domain (Figure 4E), in line with what was observed by NMR (Figure 4B). These data support the notion that the UBL and UBA domains form a stable interaction, and that the STI1 domain transiently interacts with each of the 3TH regions in the IDRs of Dsk2 (Figure 4G).

### STI1 domain and transient helices are major contributors of Dsk2 phase separatio

NMR analysis and molecular dynamics simulations revealed distinct multivalent interactions within Dsk2, including UBL-UBA, STI1-STI1, and STI1-3TH intramolecular and intermolecular interactions. We assessed the differential contributions of these interactions in Dsk2 phase separation, which depends on intermolecular interactions. We obtained phase diagrams of different domain-deletion and mutant constructs of Dsk2 (Figure 5A, 5B) and used brightfield microscopy to monitor droplet formation (at 25°C with noted protein concentration (Figure 5C)). To disrupt UBL-UBA interactions, we expressed and purified Dsk2 ΔUBL and Dsk2 mutUBA (G343A/F344A double mutant) (Waite *et al*, 2024; Sasaki *et al*, 2005). For both Dsk2 ΔUBL and Dsk2 mutUBA constructs, the loss of UBL-UBA interactions reduced phase separation propensity, as higher protein concentrations are required to induce phase separation (rightward shift in their phase diagrams compared to Dsk2 FL) (Figure 5A).

**Figure 5.**
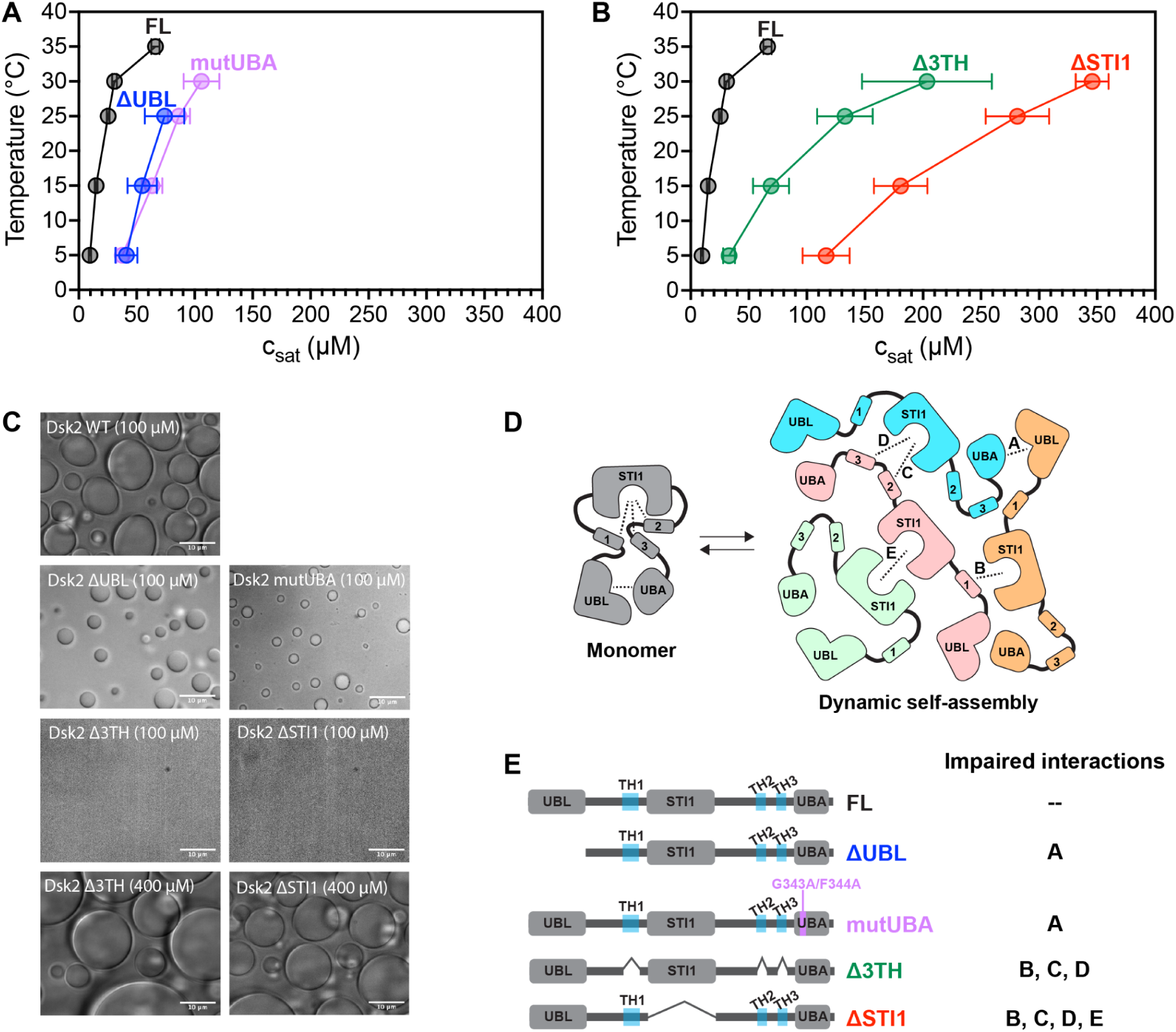
STI1 domain and transient helical regions are important drivers of Dsk2 phase separation. (A) Phase diagrams of Dsk2 ΔUBL and Dsk2 mutUBA constructs are compared against Dsk2 FL. (B) Phase diagrams of Dsk2 Δ3TH and Dsk2 ΔSTI1 are compared with Dsk2 FL. (C) Bright-field microscopy showing phase separation of Dsk2 constructs at 100 µM protein concentration (unless otherwise specified) incubated at 25°C for 20 min. Scale bars, 10 µm. (D) Representative model depicting how interactions among different regions of Dsk2 (represented by dashed lines and further described in panel E) provide the crosslinks leading to Dsk2 self-assembly and phase separation. Different monomeric units are shown in different colors where no more than two interactions among domains/regions are highlighted between the same two monomeric units for simplicity and clarity. (E) Domain architectures of Dsk2 constructs are shown, highlighting respective impaired interactions in panel D.

Removal of the three transient helical regions (Dsk2 Δ3TH) or the STI1 domain (Dsk2 ΔSTI1) resulted in much larger c_sat_ values for phase separation compared to the FL, ΔUBL, and mutUBA constructs (Figure 5B). We observed phase separation for Dsk2 Δ3TH and Dsk2 ΔSTI1, but only at substantially higher protein concentrations (Figure 5C). We propose that the pronounced reduction in phase separation propensity stems from the loss of multivalent interactions when all three transient helical regions are removed, impacting STI1-TH1, STI1-TH2, and STI1-TH3 interactions. The removal of the STI1 domain, due to eliminating STI1-STI1 interactions as well, led to the largest decrease in phase separation propensity of all constructs examined here. Given that the STI1 domain mediates multiple interactions in Dsk2 self-association, including STI1-STI1 and STI1-3TH interactions (STI1-TH1, STI1-TH2, and STI1-TH3) (Figure 5D and 5E), we conclude that the STI1 domain is the major contributor to both self-association and phase separation *in vitro*.

### Deletion of the Dsk2 STI1 domain promotes proteasome condensate formation *in vivo*

In yeast, the formation of stress-induced proteasome condensates depends on the presence of shuttle factors (Dsk2, Rad23) and polyubiquitin (Waite *et al*, 2024). Given our observations that the STI1 domain is an important driver of Dsk2 phase separation, it is likely that the STI1 domain and substrates that bind to the STI1 domain also regulate Dsk2 condensate formation in cells. We used CRISPR/Cas9 to delete the region encoding the STI1 domain of Dsk2 (*DSK2 ΔSTI1*) in wild type (WT) or *rad23Δ* cells (RAD23 deletion strain) that express GFP-tagged proteasomes (Rpn1-GFP at endogenous locus). Importantly, deletion of the STI1 domain in yeast increased the percentage of cells with at least one proteasome punctum in the *rad23Δ* genetic background under two different forms of stress, sodium azide treatment or prolonged growth (3 days) in YPD medium (Figure 6A, 6B, 6C, 6E). A slight but non-significant increase in percentage of cells with a proteasome punctum was observed in WT cells that still contained shuttle factor Rad23 when the STI1 domain of Dsk2 was removed. Additionally, the puncta appeared larger and brighter in Dsk2 ΔSTI1 cells (Figure 6A, 6B). Indeed, using a pixel intensity-based counting procedure, the percent of cells with at least one punctum exceeding a specific brightness threshold was increased in both Rad23WT and *rad23Δ* genotypes upon deletion of the STI1 domain in Dsk2 (Figure 6D, 6F).

**Figure 6.**
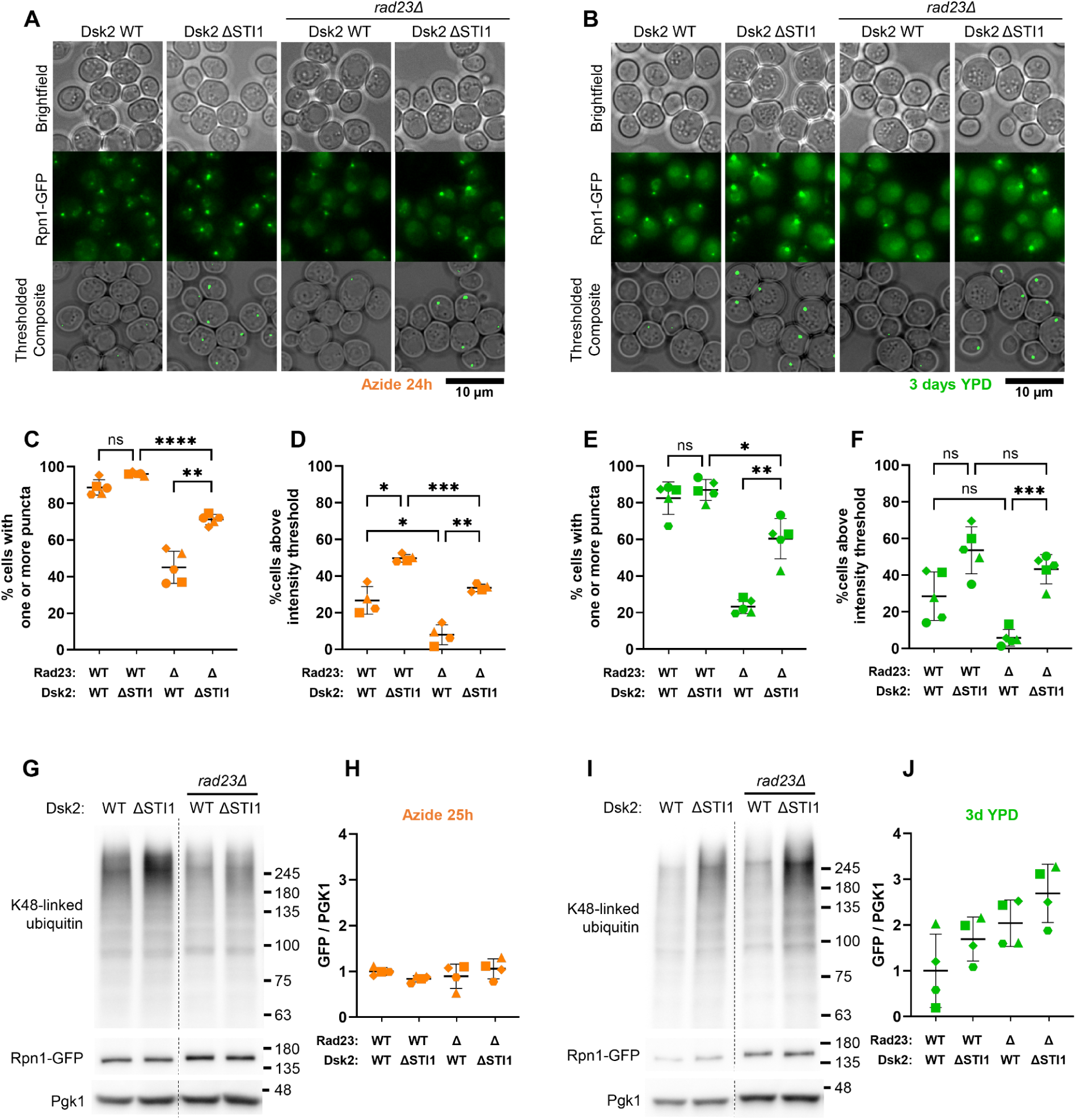
Deletion of the Dsk2 STI1 domain enhanced proteasome puncta formation. (A and B) Representative single-plane brightfield (top) and extended depth of field GFP images (middle) from five replicates for azide-treated cells (A) and cells grown for three days in YPD (3d YPD) (B). Bottom row in panels A and B display an overlay of top and middle row showing only those puncta whose brightness exceeded a defined pixel intensity threshold (used for panels D and F; see Methods). Scale bar, 10 µm. (C and E) Percent of cells with at least one punctum after 24 h azide (C) or 3d YPD (E). (D and F) Percent of cells with at least one punctum that contained at least one pixel that was brighter than the intensity threshold for azide (D) or 3d YPD (F). Respective pixel intensity thresholds were determined for each condition by setting *DSK2 ΔSTI1* to ∼50% of cells with such puncta (see Methods). (G) Western blots representative of four blots, where each lane is whole cell lysates from 0.4 OD_600_ of azide-treated cells. (H) Densitometric ratio of Rpn1-GFP (anti-GFP) to PGK1 for azide-treated cells normalized to the average ratio of WT cells. (I) and (J) are as in (G) and (H) for cells grown for 3d YPD. The means were not statistically different in H and J. For G and I, dotted lines separate two portions of a single blotted membrane. For graphs, mean ± SD is represented as black lines. Welch’s one-way ANOVA was performed for all graphs, where * P<0.05, ** P<0.01, *** P<0.001, ns not significant; ɑ=0.05. Each symbol represents one experimental replicate.

As Dsk2 contributes to the chaperone function and proteasomal degradation of many proteins, mutations impacting its function could cause changes in the amount of ubiquitinated substrates or proteasomes in the cell, consequently changing the propensity for proteasome condensate formation. Therefore, we analyzed *DSK2 ΔSTI1* strains for proteasome and polyubiquitin levels. Rpn1-GFP levels were unchanged in azide-treated cells, while there was a trend of increased Rpn1-GFP levels with deletion of the STI1 domain in cells grown for 3 days in YPD (Figure 6G-J, and Figures S7 and S8). However, there was a notable increase in K48-linked polyubiquitin levels in *DSK2 ΔSTI1* cells across both genotypes and both conditions (Figure 6G, 6I, Figures S7 and S8). Together, these data suggest that deletion of the Dsk2 STI1 domain promotes proteasome condensate formation *in vivo*, driven in part by the accumulation of polyubiquitinated proteins and, after prolonged growth, increased levels of proteasome.

## Discussion

In this work, we comprehensively mapped multivalent interactions across both folded and intrinsically-disordered domains of full-length Dsk2 on a residue-by-residue basis using NMR spectroscopy (Figure 2). No other UBQLN family member has yet been examined at this level as many resonances are significantly broadened beyond detection, specifically STI1 domains in human UBQLNs (Dao *et al*, 2018; Zheng *et al*, 2021). Among the identified multivalent interactions in Dsk2, the STI1 domain and three transient helices in disordered regions emerged as the major contributors to both Dsk2 self-association and phase separation. We further reconstituted Dsk2 condensates with colocalized proteasomes and ubiquitinated substrates, showcasing how increasing concentration of PQC components can further drive condensation of Dsk2 (Figure 1B).

Consistent with prior work on Dsk2 and human ortholog UBQLN2, our data indicate that the UBL and UBA domains interact with each other (Figure S4). The CSPs for the UBL and UBA domains (when comparing full-length to the isolated domains) map to the previously characterized UBL-UBA interface (Figure S4A-C) (Lowe *et al*, 2006). The UBL-UBA interactions occur on an intramolecular level given that Dsk2 is a monomer at low protein concentration (Figure S3). Molecular dynamics simulations of the Dsk2 monomer (where the UBL and UBA domains are unbound) resulted in a calculated effective concentration for these domains of 413.5 ± 8.8 μM (Sørensen & Kjaergaard, 2019; González-Foutel *et al*, 2022). This value is higher than the reported *K*_d_ of 80 μM for the UBL-UBA interaction using isolated domains (Lowe *et al*, 2006), further supporting that UBL and UBA domains interact intramolecularly at low protein concentrations. However, these UBL-UBA interactions become intermolecular at higher protein concentrations promoting self-association, as evidenced by the decrease in peak intensity observed in both UBL and UBA domains with increased protein concentration (Figure 3B). Unlike UBQLN2, where deletion of the UBL domain enhances phase separation (Zheng *et al*, 2021), the removal of the UBL domain in Dsk2 reduces phase separation (Figure 5A, 5C). This opposite behavior may stem from: (a) the UBL deletion in UBQLN2 also removes N-terminal disordered residues 1-33 that regulate phase separation (Dao *et al*, 2024) but this region is not present in Dsk2, (b) differences in oligomerization propensity for Dsk2 (monomer) and UBQLN2 (dimer), or (c) potentially different interaction patterns of the UBL domain with the rest of either Dsk2 or UBQLN2 (Zheng *et al*, 2021). Modulation of UBL-UBA interactions by PQC components further change Dsk2 phase separation propensity. As concentration of polyubiquitinated substrates is increased, Dsk2 phase separation is enhanced (Figure 1B). This result agrees with proteasome condensate formation in yeast, where UBL and UBA domains of Rad23 and Dsk2 contribute significantly to condensation via their interactions with proteasomes and polyubiquitin chains (Waite *et al*, 2024).

Our NMR results and molecular dynamics simulations point towards the existence of STI1-helix interactions within Dsk2 (Figure 4), which also contribute to Dsk2 self-association as protein concentration increases. Functionally, STI1 domains of co-chaperone like proteins (Sti1, HIP, SGTA, Tic40) and adaptors of the ubiquitin-proteasome system (Dsk2, UBQLNs, KPC2) are implicated in helix-mediated substrate interaction and/or activation (Fry *et al*, 2021; Schmid *et al*, 2012; Itakura *et al*, 2016; Lin *et al*, 2021). For co-chaperone Hop (an ortholog of yeast Sti1 protein), the DP2 domain (also known as STI1-II domain) directly interacts with the glucocorticoid receptor (GR) client protein via a GR amphipathic helix binding to the hydrophobic groove of the Hop DP2 domain (Figure 7A) (Wang *et al*, 2022). Similarly, recent work revealed that a transmembrane helix binds to the hydrophobic groove of the Dsk2 STI1 domain from *M. bicuspidata* (Figure 7B, S9) (Onwunma *et al*, 2024). Structural modeling using AlphaFold2 multimer predictions ((Mirdita *et al*, 2022) and Methods) suggest that the three amphipathic transient helices in Dsk2 (Figure S10A) each interact with the hydrophobic groove of the STI1 domain through their hydrophobic faces (Figure 7C-E, Figure S10B-C), reminiscent of previously characterized STI1-helix interactions in Hop and *M. bicuspidata* Dsk2 (Figure 7A-B, S9A). These AlphaFold predictions are consistent with our CSP data (Figure 4B). Additionally, STI1-3TH interactions in Dsk2 do not contribute to any significant local secondary structure stabilization in either region, as indicated by unchanged hetNOE values for 3TH regions following STI1 domain deletion (Figure 4E), or for STI1 domain following 3TH deletion (Figure S11).

**Figure 7.**
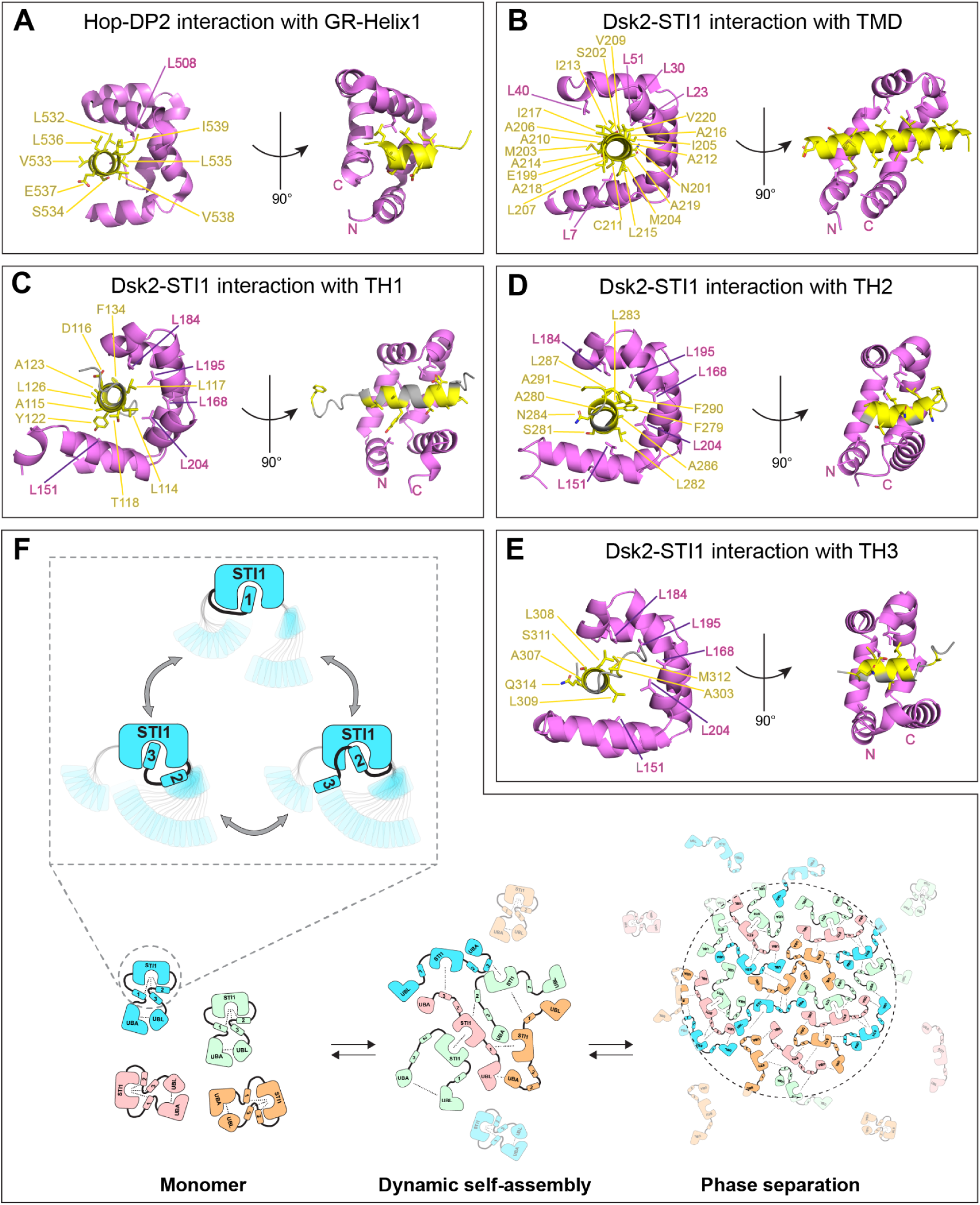
Amphipathic or hydrophobic helices bind to the hydrophobic groove of STI1 domains. (A) Structure of STI1-II (DP2) domain of Hop (pink) bound to helix-1 (yellow) of the glucocorticoid receptor client protein (PDB ID 7KW7). (B) Crystal structure of the STI1 domain of *M. bicuspidata* Dsk2 bound to a transmembrane helix (yellow) (PDB ID 9CKX); note that only relevant domains from the structures are shown in panels A-B. (C-E) Representative AlphaFold2 multimer models are shown for the interaction between the STI1 domain of Dsk2 (pink) with transient helical regions TH1 (C), TH2 (D), and TH3 (E). Only relevant short segments of transient helices are shown for clarity (TH1: residue 110-134, TH2: 278-291, TH3: 300-315), where residues with CSP > 0.04 ppm (from Figure 4B) are highlighted in yellow. See Figure S10B for full models. (F) Proposed model for how multivalent interactions in Dsk2 drive its transition from monomer to dynamic self-assembly and macromolecular phase separation. The dynamic interactions between the transient helices within the disordered regions of Dsk2 and the STI1 domain are highlighted in inset.

Recent work on the STI1 domain of Sgt2, a STI1-containing protein with chaperone function, shows that STI1 interacts with transmembrane domain (TMD) client helices with varying affinities depending on helix length and distribution of hydrophobic residues along the helix (Lin *et al*, 2021). Helices with > 11 residues containing at least 8 Leu residues were strong binders to the STI1 domain. Similarly, the three transient helices in Dsk2 vary in helix length, number and distribution of hydrophobic residues (Table S1), suggesting they may bind the STI1 domain with varying affinities. We speculate that each of these helices transiently occupies the STI1 hydrophobic groove (Figure 7), as observed in our molecular dynamics simulations (Figure 4E, 4G, Movies S1, S2). Such a mechanism of dynamic binding between STI1 and multiple helices also partially explains the weak amide resonances of the Dsk2 STI1 domain as observed by NMR spectroscopy even in Dsk2 truncation constructs (Figure S6). In the cell, we predict that the STI1-3TH interactions are likely displaced by *bona fide* STI1 substrates, as hypothesized in recent work (Onwunma *et al*, 2024). Additionally, the transient helices of Dsk2 may bind to E3 ligases to facilitate ubiquitination and proteasomal degradation of certain STI1-bound Dsk2 client proteins, as suggested for UBQLN1 interactions with mitochondrial membrane proteins (Itakura *et al*, 2016). The transient helix adjacent to the UBA domain (UBAA) in human UBQLNs interacts with the AZUL domain in the E3 ligase E6AP to potentially recruit substrates for ubiquitination (Buel *et al*, 2023).

Our solution NMR data support the existence of STI1-STI1 interactions that drive Dsk2 self-association (Figure 3, S6). In UBQLN2, we previously noted that STI1-II dimerization plays a pivotal role in driving UBQLN2 phase separation (Dao *et al*, 2018, 2024). Under identical protein concentrations and experimental conditions (50 µM, pH 6.8 buffer), Dsk2 prefers the monomeric state, unlike UBQLN2 (Figure S3). However, we do observe concentration-dependent dynamic self-assembly of Dsk2 at higher protein concentrations (Figure 3). Furthermore, a recent crystal structure of a fungal Dsk2 shows STI1 capable of forming a dimer holding two transmembrane helices in the middle (Figure S9B) (Onwunma *et al*, 2024).

Though multivalent interactions among different domains and regions potentiate self-assembly in Dsk2 (Figure 3, 5, 7G), the STI1 domain (STI1-II domain in case of UBQLN2) remains the key component driving the self-association and phase separation for both UBQLN2 (Dao *et al*, 2018) and Dsk2 (this study). The STI1 domain and its substrates are likely to regulate cellular Dsk2/proteasome condensate formation. Indeed, after deletion of the STI1 domain of Dsk2, we observed intensification of proteasome condensates in stressed yeast cells, especially when the Rad23 shuttle factor was also removed (Figure 6A, 6B). However, the effect of the absence of the STI1 domain on Dsk2/proteasome condensate formation *in vivo* was opposite to that for *in vitro* Dsk2 phase separation. The STI1 domain of Dsk2 likely binds to substrates to perform chaperone activity, and/or binds to substrates destined for degradation (Itakura *et al*, 2016; Kurlawala *et al*, 2017). Consequently, deletion of the STI1 domain may not let Dsk2 perform its chaperone activity on STI1-binding client proteins leading to accumulation of ubiquitinated substrates for degradation. Improper clearance of these substrates can lead to proteasome dysfunction and increased proteasome condensation given the propensity for ubiquitinated substrates to drive condensation (Rajendran & Castañeda, 2025; Vamadevan *et al*, 2022; Goel *et al*, 2023). Indeed, there is an increase of polyubiquitinated proteins when the STI1 domain of Dsk2 is removed (Figure 6G, 6I). Just the accumulation of polyubiquitinated substrate itself may also promote proteasome condensate formation, as we previously demonstrated for reconstituted UBQLN2/proteasome/substrate condensates (Valentino *et al*, 2024).

In summary, our work identifies and highlights the crucial role of multivalent interactions among different regions of Dsk2, particularly the STI1 domain and transient helices, in driving self-association and phase separation. Notably, interactions between the STI1 domain and transient helices closely resemble STI1-client helix interactions observed in similar STI1-containing proteins, suggesting a potential role of these transient helices in STI1’s client/substrate selectivity. The presence of different Dsk2 substrates likely modulates its inter- and/or intramolecular interactions, leading to different molecular consequences that may promote or inhibit its phase separation, both directly and indirectly, *in vitro* and *in vivo*. Interestingly, while the absence of the STI1 domain drastically reduced Dsk2 phase separation *in vitro*, we observed an apparent opposite effect on proteasome condensate formation in yeast cells *in vivo*. Putting together *in vitro* and *in vivo* observations, we propose that the Dsk2 STI1 domain, key driver of Dsk2 phase separation *in vitro*, affects condensate formation in opposing ways *in vivo*: (1) directly via STI1-STI1 or STI1-3TH interactions providing more valency for condensate formation and (2) indirectly via limiting the accumulation of polyubiquitinated proteins.

### Study Limitations

Our NMR results are collected in the absence of any additional NaCl or PEG crowding agent; these conditions do not match our *in vitro* phase separation assays. NMR experimental conditions were collected as such to improve sensitivity of NMR experiments following our previous study on UBQLN2 (Dao *et al*, 2018). For *in vivo* experiments, we could not directly assess the protein level of wild-type or modified Dsk2 due to unavailability of a suitable Dsk2 antibody for yeast lysates. Additionally, tagging Dsk2 with a C-terminal fluorescent protein drastically inhibits *in vivo* proteasome condensation formation (Waite *et al*, 2024). Stabilization or destabilization of Dsk2 resulting from the genetic or protein manipulations could affect proteasome condensate formation directly via altered protein-protein interaction landscape or indirectly via changes in proteostasis.

## Acknowledgements

We acknowledge support from National Institute of Health (NIH) R01 GM136946 to C.A.C., R35 GM149314 to J.R., R35 GM137926 to S.S., postdoctoral support to N.A. and J.K.N. from the Syracuse University Vice President of Research office, and a graduate fellowship to E.A.D. from the Madison and Lila Self Graduate Programs at the University of Kansas. S.S. acknowledges support from Alfred P. Sloan Foundation. Support for the Bruker 800 MHz spectrometer with TCI cryoprobe was provided by shared instrumentation NIH Grant 1S10OD012254. We appreciate support from Dr. Charlie Fry at the SUNY-ESF NMR facility. Additional NMR experiments were performed at the Johns Hopkins Biomolecular NMR facility with support from Dr. Ananya Majumdar. We thank Dr. Matthew Wohlever for important feedback on our work. We thank Dr. Maxwell Watkins for the SEC-MALS-SAXS experiments performed at BioCAT/SIBYLS facility. This research used resources of the Advanced Light Source (ALS), a national user facility operated by Lawrence Berkeley National Laboratory on behalf of the Department of Energy (DOE), Office of Basic Energy Sciences, through the Integrated Diffraction Analysis Technologies (IDAT) program, supported by DOE Office of Biological and Environmental Research. Additional support comes from the NIH project ALS-ENABLE (P30 GM124169) and a High-End Instrumentation Grant S10OD018483. This research also used resources of the Advanced Photon Source, a U.S. DOE Office of Science User Facility operated for the DOE Office of Science by Argonne National Laboratory under Contract No. DE-AC02-06CH11357. BioCAT was supported by grant P30 GM138395 from the National Institute of General Medical Sciences (NIGMS) of the NIH. We thank Syracuse University’s High Performance Computing cluster, OrangeZest, for computational resources and Daniel Jeski for help in running the molecular dynamics simulations. The content is solely the responsibility of the authors and does not necessarily reflect the official views of NIGMS or the NIH.

## Materials and Methods

### Protein Expression and Purification

Wild-type yeast Dsk2 was cloned from yeast genomic DNA into a pE-SUMO construct using Gibson assembly to create His-SUMO-Dsk2. Different domain deletion and mutant constructs of Dsk2 were prepared from the original plasmid using Phusion Site-Directed Mutagenesis Kit (Thermo Scientific) (Table S2). All Dsk2 constructs were expressed in *E. coli* Rosetta (DE3) cells in Luria-Bertani broth with 50 mg/L kanamycin and 35 mg/L chloramphenicol grown to OD_600_ of 0.6, induced with 0.5 mM IPTG, and expressed overnight at at 18°C for 24 h. NMR active (^15^N and ^13^C/^15^N labeled) protein samples were expressed in M9 minimal media as detailed elsewhere (Dao *et al*, 2018). Bacteria were pelleted, frozen, then lysed via freeze/thaw method in 50 mM sodium phosphate buffer pH 8.0 containing 300 mM NaCl, 25 mM imidazole, 0.5 mM EDTA, 1 mM PMSF, 1 mM MgCl_2_, and 25 U of Pierce universal nuclease. All Dsk2 constructs were purified by Ni^2+^ affinity chromatography. The N-terminal His-SUMO tag was cleaved by SUMO protease at room temperature overnight while dialyzing in a 20 mM sodium phosphate buffer, pH 7.2. To remove His-SUMO tag from the cleaved protein, the cleavage mix was passed through a Ni^2+^ or Co^2+^ column and the flow through containing purified protein was collected. To achieve a higher degree of purification, we performed anion exchange chromatography and concentrated all fractions containing purified protein using centrifugal concentrators. Protein purity was estimated by gel electrophoresis (Figure S2). Protein concentrations were measured spectroscopically using respective theoretical molar extinction coefficients (Table S3), except for Dsk2 STI1+IDR construct which lacks any Y or W residues. For Dsk2 STI1+IDR, the concentration was estimated by SDS-PAGE gel using concentration standards of similar molecular weight protein. Purified protein samples were buffer-exchanged into 20 mM sodium phosphate buffer at pH 6.8 containing 0.5 mM EDTA and 0.02% NaN_3_, and stored at -80°C.

### SEC-MALS data collection and analysis

SAXS was performed at the SIBYLS beamline (beamline 12.3.1 at the Advanced Light Source, Berkeley, CA) with in-line size exclusion chromatography (SEC) (Rosenberg *et al*, 2022; Classen *et al*, 2013; Putnam *et al*, 2007) to separate sample from aggregates and other contaminants thus ensuring optimal sample quality and multiangle light scattering (MALS) and refractive index measurement (RI) for additional biophysical characterization (SEC-MALS-SAXS). The samples were loaded on a Shodex Protein KW-803 column run by a 1260 series HPLC (Agilent Technologies) with a flow rate of 0.65 mL/min and a temperature of 25 °C. The flow passed through (in order) the UV detector (Agilent 1290 II Diode Array Detector), a MALS detector (18-angle DAWN Helios II, Wyatt Technologies), the SAXS sample cell and finally an RI detector (Optilab T-rEX, Wyatt). Scattering intensity was recorded using a Pilatus X3 2M (Dectris) detector which was placed 2.1 m from the sample giving access to a q-range of 0.011 Å-1 to 0.47 Å-1. 2.0 s exposures were acquired every 2 s during elution (∼25 minutes), and data was reduced using BioXTAS RAW 2.1.1 (Hopkins *et al*, 2017). Buffer blanks were created by averaging regions flanking the elution peak and subtracted from exposures selected from the elution peak to create the I(q) vs q curves used for subsequent analyses. Peak deconvolution by evolving factor analysis (EFA) (Meisburger *et al*, 2016) was performed in BioXTAS RAW 2.1.1. Molecular weights were calculated from the MALS and RI data using the ASTRA 7 software (Wyatt).

### Phase Diagram measurements

Protein stock samples were prepared in sodium phosphate buffer (pH 6.8, 20 mM NaPhosphate, 0.5 mM EDTA): 250-1050 µM for Dsk2 FL, Dsk2 ΔUBL, Dsk2 mutUBA, Dsk2 Δ3TH, and Dsk2 ΔSTI1. For phase separation assay, 10 µL protein sample from the stock was mixed with 10 µL of 2X phase separation buffer (pH 6.8, 20 mM NaPhosphate, 300 mM NaCl, 15% PEG8K, 0.5 mM EDTA) and incubated at different/desired temperatures for 20 min, followed by centrifugation at 15000 g for 5 min at respective temperature. After centrifugation, 10 µL supernatant was immediately pipetted out without disrupting the pellet and mixed 1:1 in 8 M urea solution to quench any further phase separation due to temperature changes. The concentration of the dilute phase representing the saturation concentration or c_sat_ (1:1 urea diluted supernatant) was measured spectroscopically on NanoDrop One (Thermo Scientific) using respective molar extinction coefficients (c_sat_ = (A280/molar extinction coefficient) *2). Phase separation assays for each Dsk2 construct were performed with two or more protein preparations in triplicates.

### Microscopy

For Dsk2 and Ub-substrate concentration dependent phase separation study (Figure 1B), samples were prepared to contain 10 μM Dsk2 (spiked with 20 nM Dsk2 labeled with Alexa Fluor 647) and 0-200 nM of K63-linked ubiquitinated substrates (Rsp5-ubiquitinated R-Neh2Dual-sGFP) in 50 mM Tris-Cl pH 7.5 buffer containing 5 mM MgCl2, 5% glycerol, 1 mM ATP, 10 mM creatine phosphate, 0.1 mg/mL creatine phosphokinase, 2 mM DTT, 1 mg/mL BSA, and 1% DMSO, 3% PEG 8000. K63-linked ubiquitinated substrate was prepared as described in (Valentino *et al*, 2024). Substrate was ubiquitinated in a 500 µL reaction then purified via size exclusion chromatography on a Superdex S200 column; fractions with the greatest extent of ubiquitination were pooled and concentrated using centrifugal concentrators. For samples containing proteasome in addition to Dsk2 and Ub-substrate (Figure 1C), 100 nM TagRFP-T-Rpn6 containing proteasome (as described in (Valentino *et al*, 2024)) were added in above condition with 100 µM proteasome inhibitor cocktail. For bright-field imaging of phase separation (Figure 1D, Figure 5C), 50 µL samples were prepared as described for the phase separation assay from 200 µM protein stocks for all Dsk2 constructs. For Dsk2 Δ3TH and Dsk2 ΔSTI1, additional samples were prepared from 800 µM protein stock.

All samples were added on to Eisco Labs Microscope Slides, with single Concavity, and covered with 5% BSA coated coverslips to minimize potential droplet-surface interactions. Sample slides were incubated coverslip-side down at 25°C for 20 min (or 18°C for 1 h: Figure 1B and 1C) prior to imaging. All samples were imaged on an ONI Nanoimager (Oxford Nanoimaging Ltd) equipped with a Hamamatsu sCMOS ORCA flash 4.0 V3 camera using an Olympus 100X/1.4 N.A. objective. Images were prepared using Fiji (Schindelin *et al*, 2012) and (Mutterer & Zinck, 2013).

### NMR experiments

All NMR experiments were performed at 25**°**C on a Bruker Avance III 800 MHz spectrometer equipped with TCI cryoprobe. Proteins were prepared in the NMR buffer: 20mM sodium phosphate buffer (pH 6.8), 0.5mM EDTA, 0.02% NaN_3_, and 5% D_2_O. All NMR data were processed on NMRBox (Maciejewski *et al*, 2017) using NMRPipe (Delaglio *et al*, 1995) and analyzed using CCPNMR 2.5.2 (Vranken *et al*, 2005).

### NMR spectra

^1^H-^15^N TROSY-HSQC experiments were acquired using spectral widths of 15 and 27 ppm in the direct ^1^H and indirect ^15^N dimensions, and corresponding acquisition times of 200 ms and 46 ms. Centers of frequency axes were ∼4.7 and 117.5 ppm for ^1^H and ^15^N dimensions, respectively. ^1^H-^15^N TROSY spectra were processed and apodized using a Lorentz-to-Gauss window function with 15 Hz line sharpening and 20 Hz line broadening in the ^1^H dimension, while ^15^N dimension was processed using a cosine squared bell function. Chemical shift perturbations (CSPs) were quantified as follows: Δδ = [(ΔδH)^2^ + (ΔδN/5)^2^]^1/2^ where ΔδH and ΔδN are the differences in ^1^H and ^15^N chemical shifts, respectively.

### NMR chemical shift assignments

We determined backbone ^1^H, ^13^C, ^15^N assignments using traditional ^1^H-detect triple-resonance experiments (HN(CA)N, HN(COCA)N, HNCACB, CBCA(CO)NH) on ^13^C/^15^N samples containing 250 µM Dsk2 FL or 400 µM Dsk2 UBL or 180 µM Dsk2 UBA or 400 µM Dsk2 ΔSTI1 proteins in pH 6.8 NMR buffer at 25°C. All experiments used optimized parameter sets incorporating non-uniform sampling. Assignments for the STI1 domain were obtained using a 100 µM sample of the Dsk2 IDR+STI1 construct (residues 76-223, inclusive of the IDR between the UBL and STI1, and the STI1 domain) with standard triple resonance experiments (HN(CA)N, HNCACB, CBCA(CO)NH, HNCO, and HN(CA)CO). Approximately 88% of backbone amide assignments were transferred by visual inspection to full-length Dsk2 NMR spectra.

Acquisition times for ^1^H-detect experiments were 20 ms, 16 ms, 6 ms, and 75-80 ms, in the indirect ^15^N dimensions, indirect ^13^CO, indirect ^13^Cα/Cβ dimensions, and direct ^1^H dimensions, respectively. Spectral widths were generally 16 ppm in indirect ^13^CO (for ^1^H-detect experiments), 28 ppm in indirect ^15^N (for ^1^H-detect experiments), and 70 ppm in indirect ^13^C_α_/C_β_. Experiments were acquired with 12%–25% sampling using the Poisson Gap sampling method (Hyberts *et al*, 2010). Spectra were processed using NMRPipe and employed standard apodization parameters and linear prediction in the indirect dimensions, and analyzed using CCPNMR on NMRBox. Using these experiments, we successfully assigned backbone resonances (H, N, C_α_, C_β_, CO) for ∼84% of all residues.

### Secondary Structure Determination from NMR Data

For C_α_ and C_β_ secondary shift calculations, random coil chemical shifts for Dsk2 constructs were determined at https://spin.niddk.nih.gov/bax/nmrserver/Poulsen_rc_CS/ using default parameters at 25°C sample temperature and pH 6.8 (Kjaergaard & Poulsen, 2011). We calculated ΔδCα values (Figure 2B top) by combining C_α_ secondary shifts from several Dsk2 constructs (Dsk2 FL, Dsk2 UBL only, Dsk2 UBA only, Dsk2 IDR+STI1). The predicted C_α_ chemical shifts of Dsk2, presented in Figure 2B bottom, were obtained using the EFG-CS web server (https://biosig.lab.uq.edu.au/efg_cs/) (Gu *et al*, 2024). This web server employs machine learning-based models to predict chemical shifts from protein sequences and structures, including experimental structures and AlphaFold2 predictions. For this study, we used the AlphaFold2-predicted structure of Dsk2 (AF-P48510-F1-v4) as input to generate the predicted C_α_ chemical shifts.

### 15N Relaxation experiments

Longitudinal (R_1_), transverse (R_2_) backbone ^15^N relaxation rates, and {^1^H}-^15^N steady-state heteronuclear Overhauser enhancement (hetNOE) were measured for Dsk2 and its domain deletion variants using established interleaved relaxation experiments and protocols (Castañeda *et al*, 2016; Hall & Fushman, 2003). Protein concentrations used were either 50 µM or 400 µM. Relaxation inversion recovery periods for R_1_ experiments were 4 ms (x 2), 600 ms (x 2), and 1000 ms (x 2) for Dsk2 FL and 4 ms (x 2), 500 ms (x 2), and 800 ms (x 2) for Dsk2 ΔSTI1, using an interscan delay of 2.5 s. Total spin-echo durations for R2 experiments were 8 ms (x 2), 32 ms, 48 ms, 64 ms (x 2), 120 ms, and 160 ms (x 2) for Dsk2 FL and Dsk2 ΔSTI1, and 4 ms (x 2), 32 ms (x2), 48 ms, 64 ms (x 2), 96 ms (x2), and 120 ms (x 2) for Dsk2 Δ3TH using an interscan delay of 2.5 s. Heteronuclear NOE experiments were acquired with an interscan delay of 4.5 s. All relaxation experiments were acquired using spectral widths of 12 and 28 ppm in the ^1^H and ^15^N dimensions, respectively, with corresponding acquisition times of 90 ms (or 75 ms for 50 µM ΔSTI1 R_1_) and 28 ms (or 26 ms for 50µM ΔSTI1 R_1_). Spectra were processed using squared cosine bell apodization in both ^1^H and ^15^N dimensions. Relaxation rates were derived by fitting peak heights to a mono-exponential decay using RELAXFIT (Fushman *et al*, 1997). Errors in R_1_ and R_2_ were determined using 500 Monte Carlo trials using RELAXFIT on MATLAB. Errors in hetNOE measurements were determined using the standard error (SE) propagation formula.

### CALVADOS Molecular Dynamics Simulations

We investigated the interactions of the transient helical (TH) regions with the STI1 groove by performing single chain simulations of full-length Dsk2 using CALVADOS3_COM_ (Tesei *et al*, 2021; Cao *et al*, 2024; Tesei & Lindorff-Larsen, 2023). In CALVADOS3_COM_, each residue is represented as a single bead placed at the center of mass calculated from all atoms within the residue (Cao *et al*, 2024). All interactions were assigned as described, and the elastic network model was applied to the folded domains to restrain non-bonded pairs (UBL: residues 1-75, STI1: residues 147-223, and UBA: residues 327-373). Ten full-atom initial conformations of Dsk2 were generated using Modeller (Šali & Blundell, 1993) from the AlphaFold predicted structure

(AF-P48510-F1-v4), which were then mapped to the CALVADOS3_COM_ coarse-grained representation. Simulations were run at a temperature of 298.15K, pH of 6.8, and ionic strength of 0.22 M for 70 ns, where the first 3.5 ns was discarded as equilibration, using Langevin dynamics. The drag coefficient was set to 0.01 ps^-1^, and the timestep was set to 10 fs (Cao *et al*, 2024).

A second set of simulations were performed for full-length Dsk2 where the UBL and UBA domains were placed in a bound conformation following the crystal structure PDB ID 2BWE (Lowe *et al*, 2006). To generate a starting structure, the AlphaFold predicted structure was modified in Pymol such that the UBA was moved to an orientation overlapping that of PDB ID 2BWE. The 10 amino acids (residues 317-326) leading to the UBA domain were rebuilt using the Build tool within Pymol, before being energy-minimized using Relax_Amber (Mirdita *et al*, 2022). Modeller (Šali & Blundell, 1993) was then used to generate ten all-atom initial conformations. To ensure that the UBL and UBA remained bound within the simulations, the elastic bond network was applied to UBL and UBA as a group, and then applied to STI1 separately. Simulations were then performed following the same methodology as the unbound Dsk2 structure.

To quantify the occupancy of the STI1 groove, the amino acids of STI1 were labeled as either interior (being within the hydrophobic groove) or exterior. The center position of the STI1 groove was calculated by averaging the positions of the interior residues. Each IDR residue was then compared to the center position of STI1, and if it was closer to the center position than the furthest interior residue, the residue was initially considered to be within the groove. For residues marked as within the groove, we further verified occupancy by checking that the residue was closer to an interior residue than an exterior residue. Lastly, to account for the groove’s curvature, a principal component analysis was performed on the interior group to identify the dominant axis of the groove, and marked residues as within the groove that were similar to this axis. The presented probabilities are the averages from 10 independent simulations, with each trajectory being the probability of the IDR occupying the STI1 groove within the 133,000 frames of each trajectory. All trajectory data is available at (https://zenodo.org/records/15333863) and analysis code is available at https://github.com/sukeniklab/Dsk2_transient_helices.

### AlphaFold2 multimer modeling for predicting interactions between the STI1 domain and transient helices of Dsk2

We used AlphaFold2 Multimer (ColabFold v1.5.5) (Mirdita *et al*, 2022) to model how the STI1 domain interacts with transient helices in Dsk2. We used the following residues of Dsk2 FL as input: 147-228 as the STI1 domain, 77-146 as TH1 (this is inclusive of the IDR between the UBL domain and the STI1 domain), 229-291 as TH2 (this is inclusive of the IDR between the STI1 domain and TH2 region), and 292-325 as TH3. The exact amino acid sequence for these regions are detailed in Table S2. Each multimer run resulted in five predicted models ranked 1 to 5. Overlays of all five models from each multimer run are shown in Figure S10B. For simplified representation, only one model from each multimer run is shown in Figure 7C-E and S10C (rank 2 for TH1; rank 3 for TH2; rank 1 for TH3 – each chosen at random).

### Yeast strains and gene manipulations

*S. cerevisiae* strains used in this work are reported in SI Table 5. All strains harbor Rpn1 C-terminally tagged with eGFP (“Rpn1-GFP”) (Waite *et al*, 2024). Strains with Rad23 and/or Dsk2 deletion and replacement with antibiotic resistance genes were also generated previously using standard PCR-based procedures (Waite *et al*, 2024; Janke *et al*, 2004; Goldstein & McCusker, 1999). Strains with deletion of the STI1 domain were obtained using CRISPR-Cas9. Candidate guide RNAs were determined using the CRISPR/Cas9 target online predictor CCTop (Stemmer *et al*, 2015; Labuhn *et al*, 2018). The sequence coding for the selected guide RNA was cloned into plasmids derived from 2µ CRISPR vector pML107 (Addgene plasmid # 67639, (Laughery *et al*, 2015)), reported in Table S5. The sgRNA/CRISPR plasmid and the repair duplex were transformed into yeast and mutations were confirmed by sequencing the genomic region. Yeast stocks of cells that no longer harbored the plasmid were used in experiments.

### Yeast growth conditions

*S. cerevisiae* yeast were grown at 30°C in liquid yeast extract peptone medium with 2% dextrose (YPD) with shaking or on YPD agar plates. Prior to experimentation, yeast strains were streaked from YPD-glycerol stocks onto YPD agar plates, which were either stored at 22-30°C for up to 4 days or stored for an additional two to three weeks in a refrigerator (∼4°C). Cells were inoculated from one or two isolated colonies on plates into liquid YPD medium and grown overnight to 20 to 24 OD_600_. Cultures were diluted to ∼0.5 OD_600_ (mean 0.54) in fresh YPD media and grown for three days (72–76 h). For azide treatment, cells inoculated at ∼0.5 OD_600_ in fresh YPD were grown for 4 h before sodium azide was added to a final concentration of 0.5 mM for an additional 24–25 h (24 h azide). After three days in YPD, the optical densities were 43–52 OD_600_; after 25–26 hours in azide, the optical densities were 4.7–8.9 OD_600_. These optical densities were used to collect cells for whole cell lysis.

### Yeast live cell microscopy

Live yeast cells were diluted in PBS buffer (200 µL culture + 800 µL of PBS) to reduce autofluorescence from the YPD. Cells were pelleted by centrifugation (2,000 x g, 2 min, room temperature) and resuspended in approximately 5–10 µL (azide) or approximately 50 µL (YPD) of the supernatant after aspiration of most of the supernatant to obtain a cell density appropriate for capture of at least 200 cells across one or two fields of view. Three microliters of the resuspended cells were immobilized on a microscopy slide using an agarose gel pad (1% in PBS) that was approximately 0.2 mm thick (modified from https://www.youtube.com/watch?v=ZrZVbFg9NE8, 2019). Images were acquired at room temperature on a Keyence BZ-X810 microscope using a Nikon Plan Apo 100X/1.40 objective and the onboard 2.8 megapixel monochrome CCD camera. Brightfield images were captured as single-plane images. Two-dimensional GFP images of whole cells were captured with the Keyence BZ-X GFP filter cube (Ex 470/40 nm; Em 525/50 nm) using the Quick Full Focus (extended depth of field) setting in the BZ-X800 Viewer application. This setting captures in-focus light in steps of ∼0.8 µm into a two-dimensional 8-bit image. Exposure time was generally 1.2 seconds per step. Images of two non-overlapping fields of view of a sample on a single agarose pad, at least 400 µm apart, were captured within 10 minutes of dilution of cells in PBS, to avoid dissipation of granules that occurs after extended time.

### Quantification of proteasome puncta

To count the percent of cells with proteasome puncta, >212 cells per experimental replicate per strain (min. 212, mean 362 ± 94 (s.d.), max. 710) were manually counted in randomized, blinded images to determine the percent of green-fluorescing cells that have at least one punctum. We observed that the deletion of the Dsk2 STI1 domain made proteasome puncta appear brighter. To support this qualitative observation with an intensity-based metric, Fiji (Schindelin *et al*, 2012) was used to count cells with proteasome puncta whose brightness exceeded a defined pixel intensity threshold as follows. The grayvalue threshold was defined so that around 50% of the Dsk2-ΔSTI1 cells in one unblinded test image for each treatment condition contained at least one pixel from an Rpn1-GFP punctum that met or exceeded the grayvalue. For azide-treated cells, the threshold was at a gray value of 25 (8-bit image, 256 total gray levels; Figure 6D). For cells grown for three days in YPD, the ∼50% threshold was 35 (Figure 6F). This ∼50% threshold definition was chosen to avoid inclusion of pixels that did not correspond to proteasome puncta (e.g. brighter pixels in the nucleus) while providing a suitable range of measurements between strains that have fewer and dimmer puncta and cells with more and brighter puncta. The threshold excluded diffuse nuclear and cytoplasmic signals. A mask was created from the thresholded pixels, and was overlaid on the corresponding brightfield image (as in Figure 6). ImageJ macros in Fiji were used to perform the pixel thresholding, masking, and composite image generation (Figure S12). Composite images were randomized and blinded before cells were manually counted with the Cell Counter tool.

### Cell lysis and immunoblotting analysis

At 25–26h after azide treatment or at 75–76 hours of culture in YPD (3d YPD), the equivalent of two OD_600_ cell amount was harvested by centrifugation and immediately lysed into sample buffer and frozen at −80 °C as follows. Alkaline lysis was carried out as previously reported (Kushnirov, 2000). In short, pellets were resuspended in 100 μL deionized water to which 100 μL 200 mM NaOH was added, and samples were incubated at room temperature for 5 min. Cell suspensions were pelleted, resuspended in 50 μL SDS-PAGE sample buffer (0.06 M Tris–HCl, pH 6.8, 5% glycerol, 2% SDS, 4% β-mercaptoethanol, 0.0025% bromophenol blue), boiled at 98 °C for 5 min, and supernatant was collected and frozen at −80 °C. SDS-PAGE of 10 µL (∼0.4 OD) samples was followed by western blotting. Membranes were analyzed by immunoblotting using antibodies against GFP (Chromotek, catalog no. 3h9), PGK1 (Invitrogen, catalog no. 459250), Rad23 (R&D Systems, catalog no. AF4555), and K48-linked polyubiquitin (Cell Signaling Technology, catalog no. 8081). Images were acquired using a G:Box imaging system (Syngene) and captured with GeneSys software. Statistical analyses were performed in GraphPad Prism.

