## Supplementary Information for "STI1 domain dynamically engages transient helices in disordered regions to drive self-association and phase separation of yeast ubiquilin Dsk2"

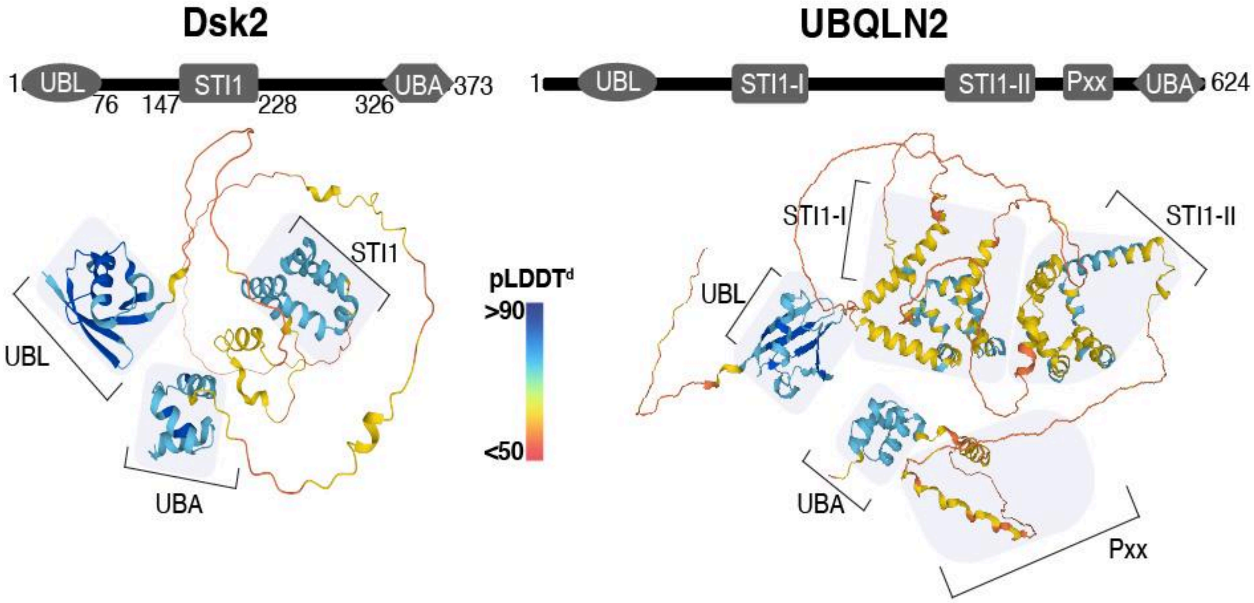

**Figure S1. Structural (and/or domain) similarities in Dsk2 and UBQLN2.** AlphaFold predicted structures of Dsk2 and UBQLN2 are compared along with their respective domain architectures. Individual domains have been labeled. Protein secondary structures have been colored according to their prediction confidence score (pLDDT score).

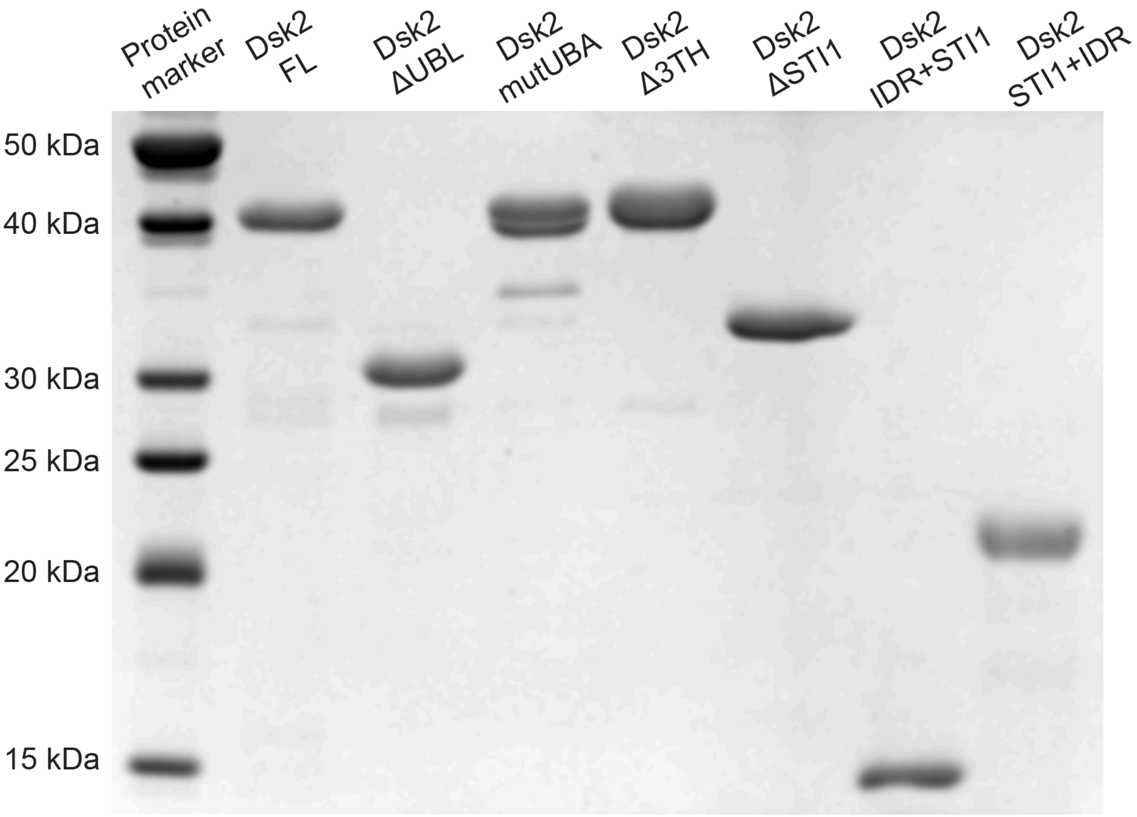

**Figure S2. Dsk2 expression.** SDS-PAGE analysis shows that all Dsk2 constructs (Dsk2 FL, Dsk2 ΔUBL, Dsk2 mutUBA, Dsk2 Δ3TH, Dsk2 ΔSTI1, Dsk2 IDR+STI1, Dsk2 STI1+IDR) were successfully expressed and purified with high purity (>90%).

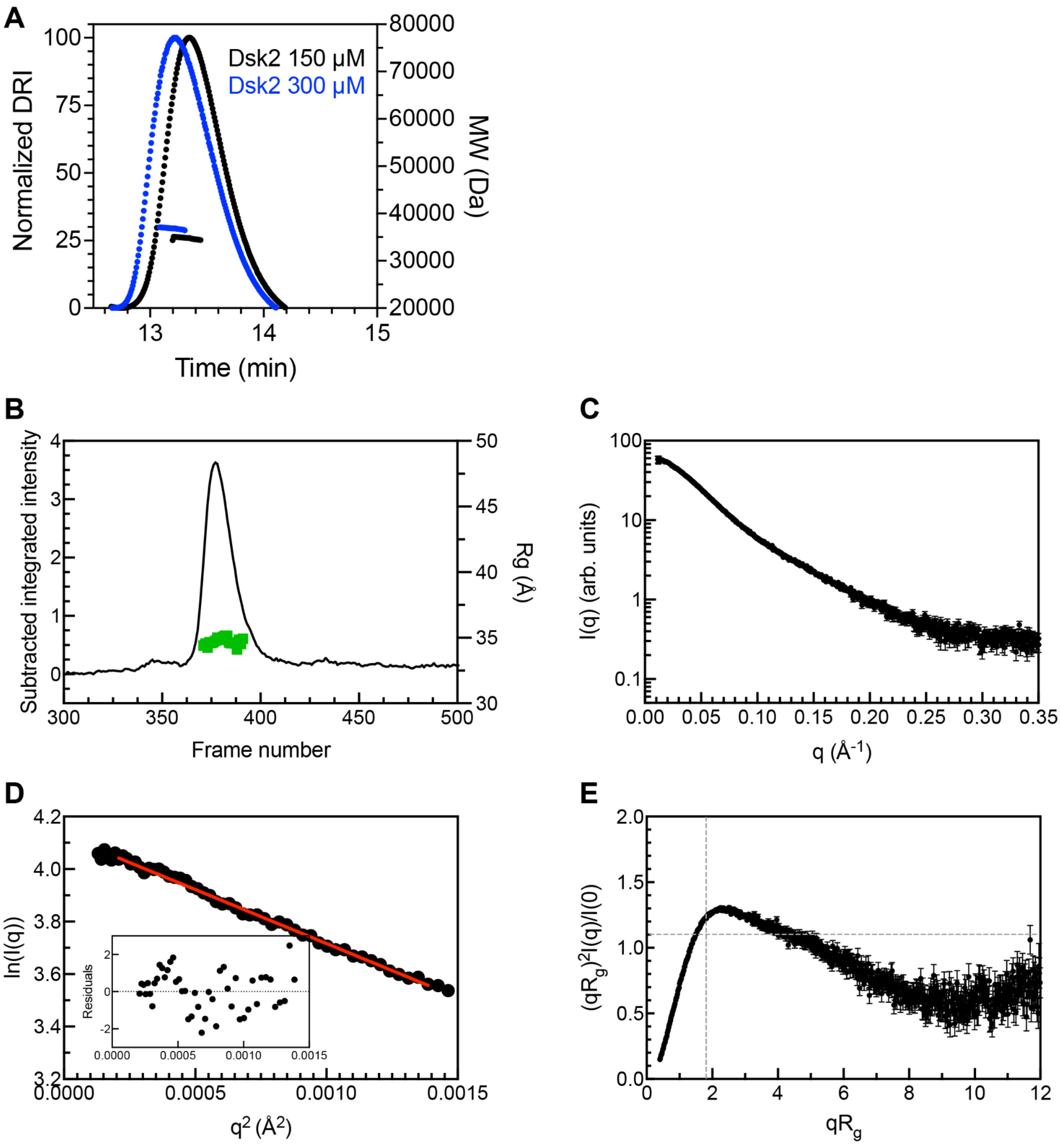

**Figure S3. Dsk2 predominantly exists in a monomeric state**. (A) SEC-MALS analysis shows that Dsk2 is a monomer at 150 μM (black) and 300 μM (blue) protein concentration using NMR buffer at pH 6.8. A slightly earlier elution at the higher concentration suggests formation of dynamic higher order self-assemblies/association. (B) SEC-SAXS profiles for Dsk2 at 150 μM with green denoting radius of gyration ($R_g$) values on the right y-axis. (C) I(q) vs. q scattering curve determined from frames 379-390 on the corresponding SEC-SAXS profiles in B. (D) Guinier plot with red line showing the linear fit of ln(I(q)) vs. $q^2$, while inset shows residuals of fit. (E) Dimensionless Kratky plot includes dashed lines to indicate where a globular protein would peak.

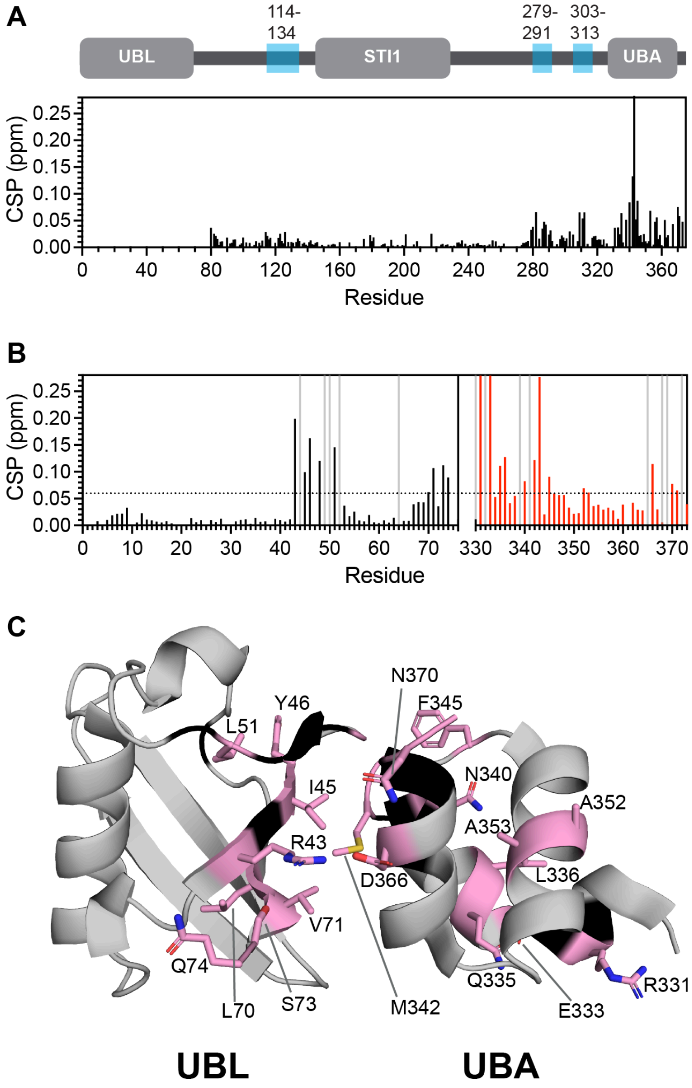

**Figure S4. The UBL domain of Dsk2 interacts with its UBA domain.** (A) Residue-level CSPs between Dsk2 FL and Dsk2 ΔUBL show large CSPs for residues in the UBA domain of Dsk2 indicative of UBL-UBA interactions. (B) CSPs on a per-residue basis are shown between Dsk2 UBL-only and Dsk2 FL (black bars), and between Dsk2 UBA-only and Dsk2 FL (red bars). Gray bars represent residues for which no amide peaks were observed (or unassigned) for Dsk2 FL. (C) Residues with CSPs > 0.06 ppm (above dotted black line in panel B) are highlighted as pink sticks and map to the UBL:UBA interface as shown on the crystal structure of the bound form of isolated Dsk2 UBL and UBA domains (PDB: 2BWE).

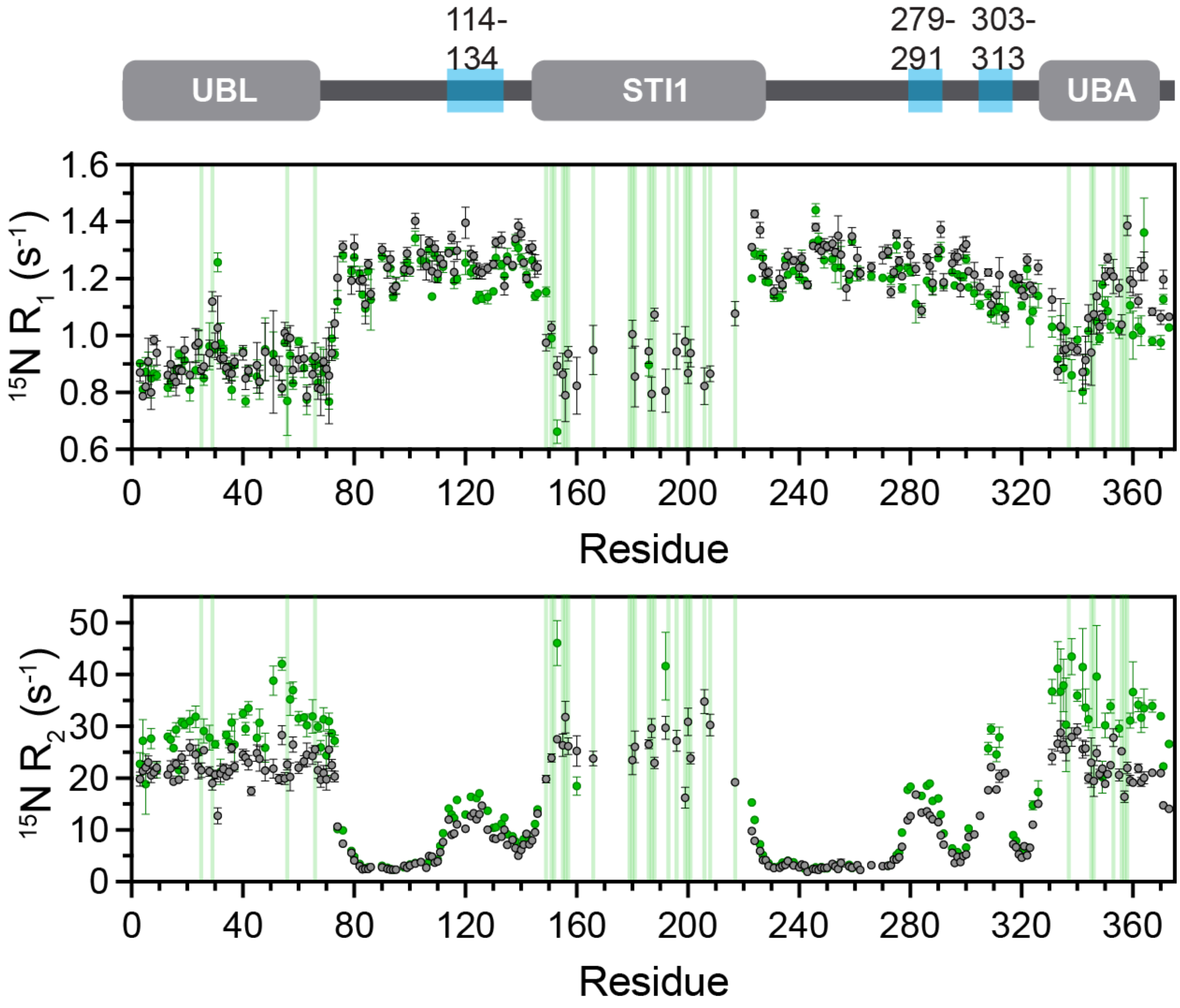

**Figure S5. Full-length Dsk2 $^{15}N$ $R_1$ and $R_2$ relaxation rates are concentration-dependent.** $^{15}N$ $R_1$, and $R_2$ relaxation rates are compared for Dsk2 at 50 μM (gray) and 400 μM (green). Errors in relaxation rates were determined using 500 Monte Carlo trials using RELAXFIT (see Methods). The concentration-dependent increase in $R_2$ relaxation rates (and corresponding decrease in $R_1$ relaxation rates) for resonances in the UBL and UBA domains suggest increased UBL:UBA intermolecular interactions with increased protein concentration. Green bars represent resonances for which there is no observable amide resonance at 400 μM; primarily affected are resonances corresponding to the STI1 domain, suggestive of intermolecular interactions involving the STI1 domain.

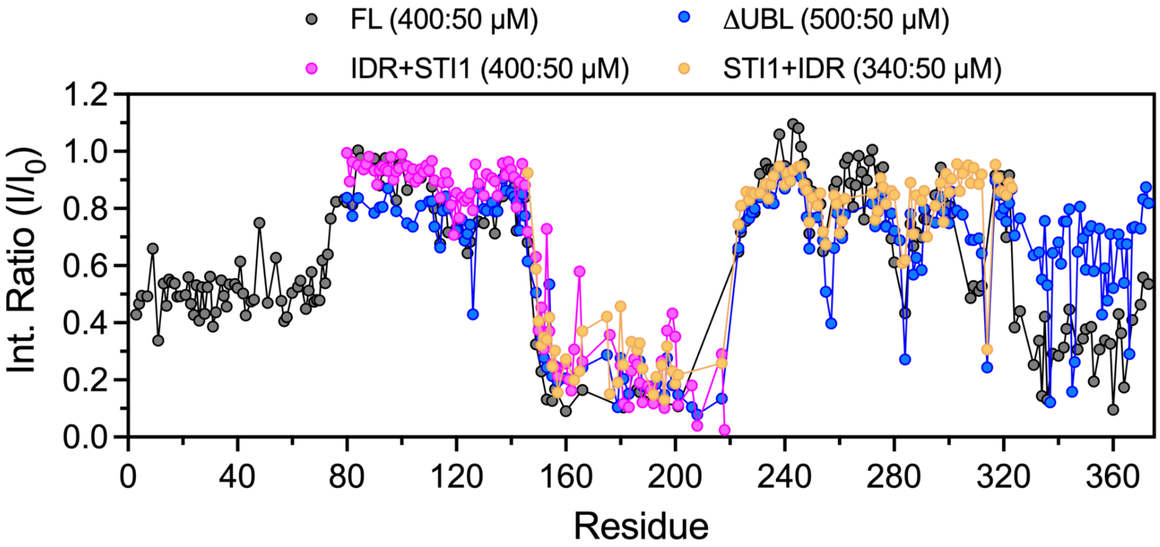

**Figure S6. STI1-STI1 interactions indicative of self-association of Dsk2 variants.** Residue-specific concentration-dependent intensity ratios ($I/I_0$) are plotted between low ($I_0$) and high (I) protein concentrations of STI1 domain-containing Dsk2 variants (Dsk2 FL, Dsk2 dUBL, Dsk2 IDR+STI1, and Dsk2 STI1+IDR). Concentrations are noted in the legend above plot; intensity ratio is corrected for differences in protein concentration and number of scans used (see Methods). Notably, residues within the STI1 domain of <u>all</u> Dsk2 constructs exhibit a similar decrease in peak intensities at higher protein concentration indicative of concentration-dependent STI1-STI1 interactions.

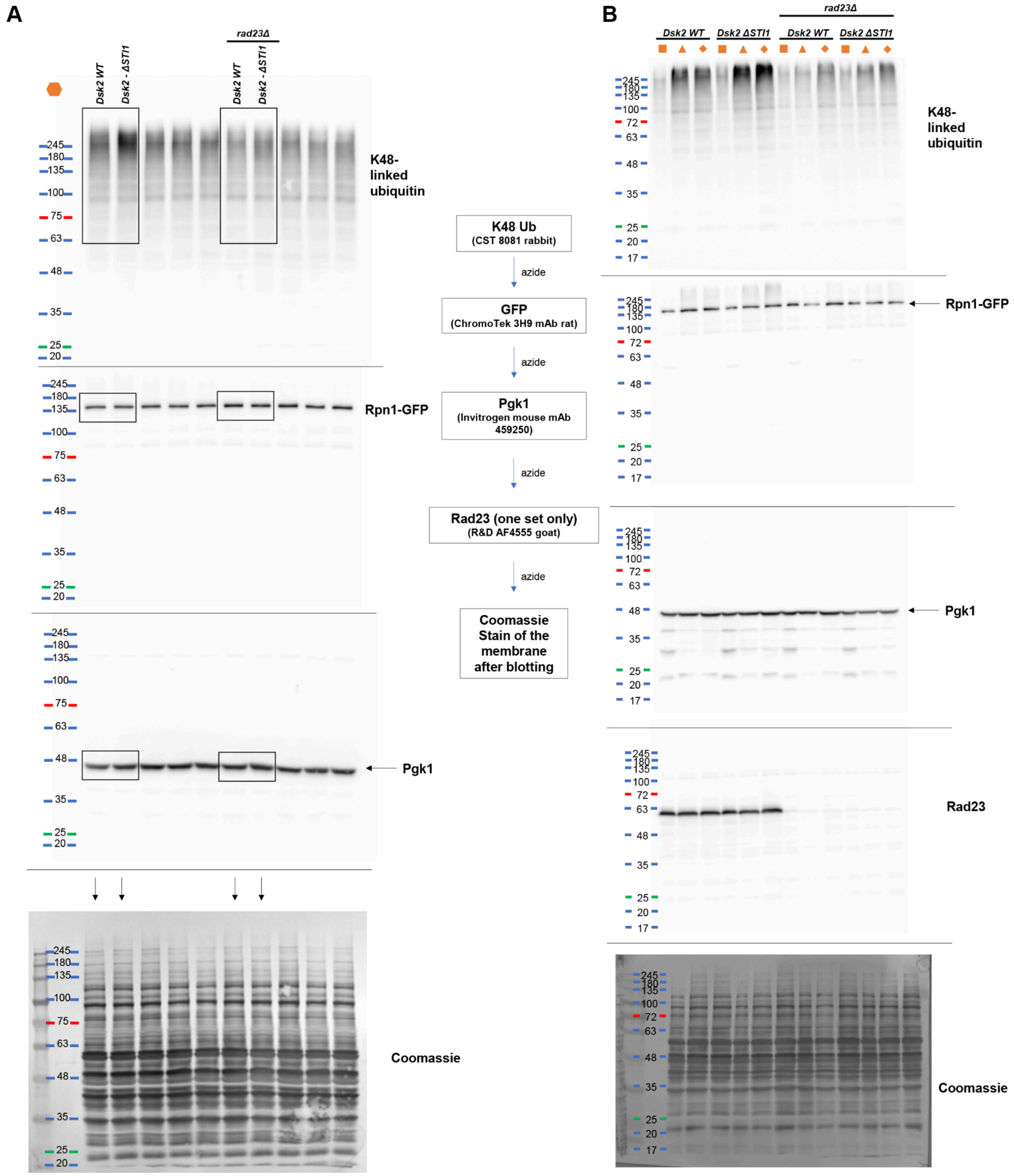

**Figure S7. Full western blots for azide-treated yeast cell lysates.** (A) Full membranes of the western blots depicted in Figure 6G (highlighted in boxes). (B) Full membranes with blots from lysates of the other three replicates quantified in Figure 6H. Selected lanes from panel A (highlighted in boxes) and all lanes from panel B were included in the GFP/PGK1 quantification. 0.02% sodium azide in TBST was used to inactivate the horseradish peroxidase enzyme conjugated to the respective secondary antibodies between reprobes of the same membranes with different primary antibodies, which were from different species (in the order as shown in the middle flowchart). The symbols in panels A and B represent individual replicates and correspond to the symbols in the graphs of Figure 6.

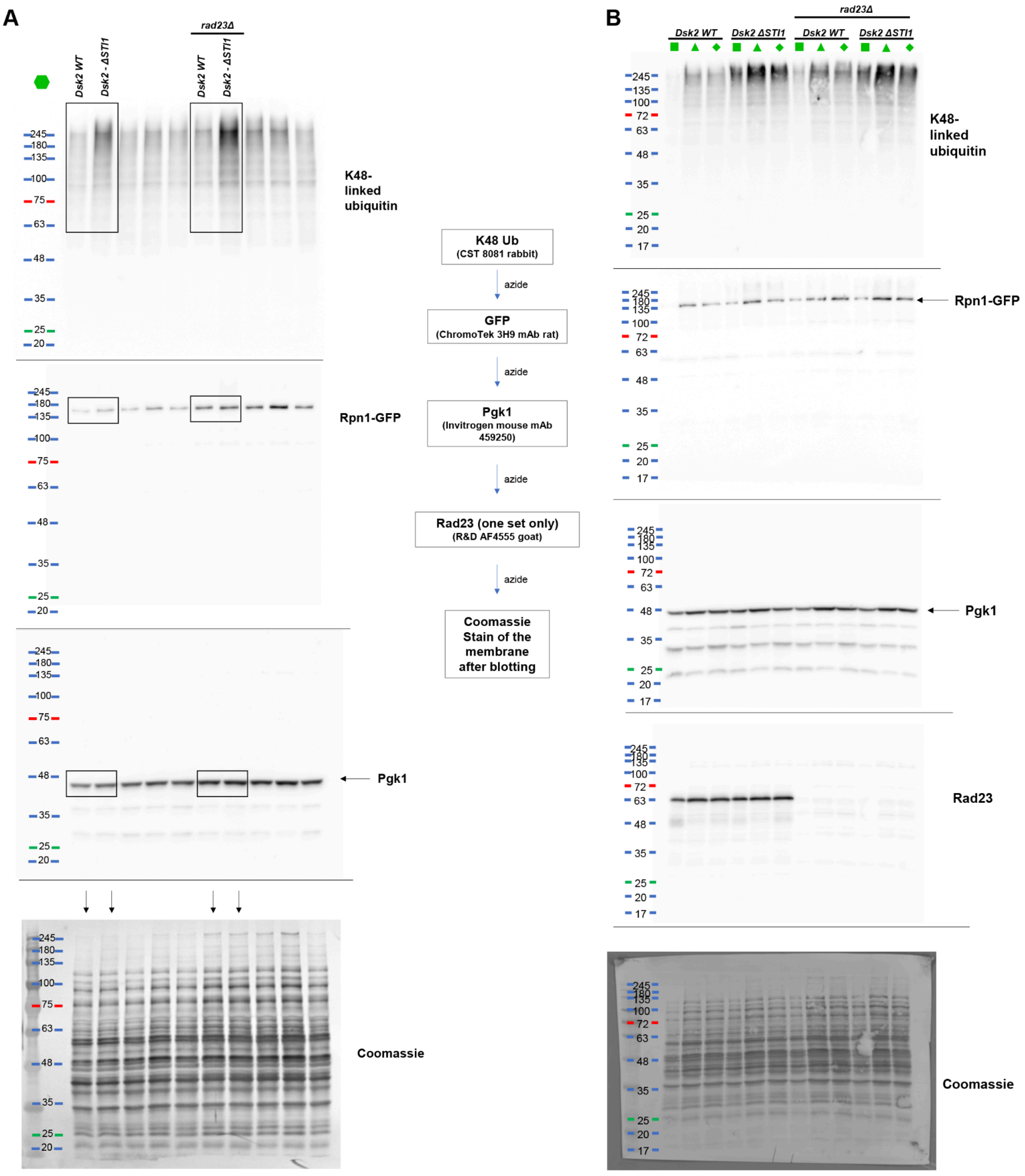

**Figure S8. Full western blots for 3d YPD (cells grown for 3 days in YPD) yeast cell lysates.** (A) Full membranes of the western blots depicted in Figure 6I (highlighted in boxes). (B) Full membranes with blots from lysates of the other three replicates quantified in Figure 6J. Selected lanes from panel A (highlighted in boxes) and all lanes from panel B were included in the GFP/PGK1 quantification. 0.02% sodium azide in TBST was used to inactivate the horseradish peroxidase enzyme conjugated to the respective secondary antibodies between reprobes of the same membranes with different primary antibodies, which were from different species (in the order as shown in the middle flowchart). The symbols in panels A and B represent individual replicates and correspond to the symbols in the graphs of Figure 6.

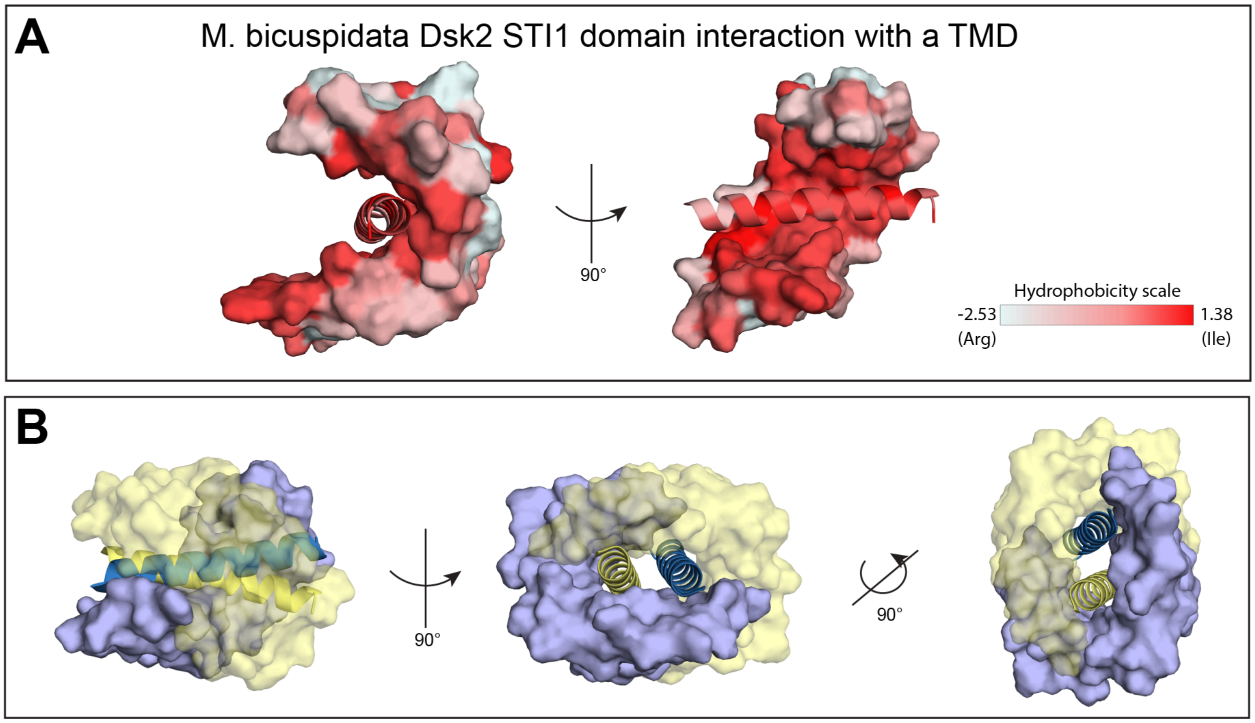

**Figure S9. Hydrophobic interactions between the *M. bicuspidata* Dsk2 STI1 domain and TMD.** (A) Surface representation of *M. bicuspidata* Dsk2 STI1 domain's hydrophobic grove bound to a hydrophobic transmembrane domain (TMD) helix (cartoon representation) (PDB 9CKX) for only one of the bound STI1-TMD (in *trans* configuration). All amino acid residues are colored from white to red based on their increasing hydrophobicity using Pymol. (B) A structural representation of the STI1-TMD chimeric protein dimer (PDB 9CKX) highlighting STI1-TMD interactions in trans configuration that mediates dimerization (Onwunma et al, 2024). Individual STI1 domains are shown as blue and yellow surfaces, while individual TMD helices attached to respective chimeric molecules are shown in blue and yellow cartoon representations.

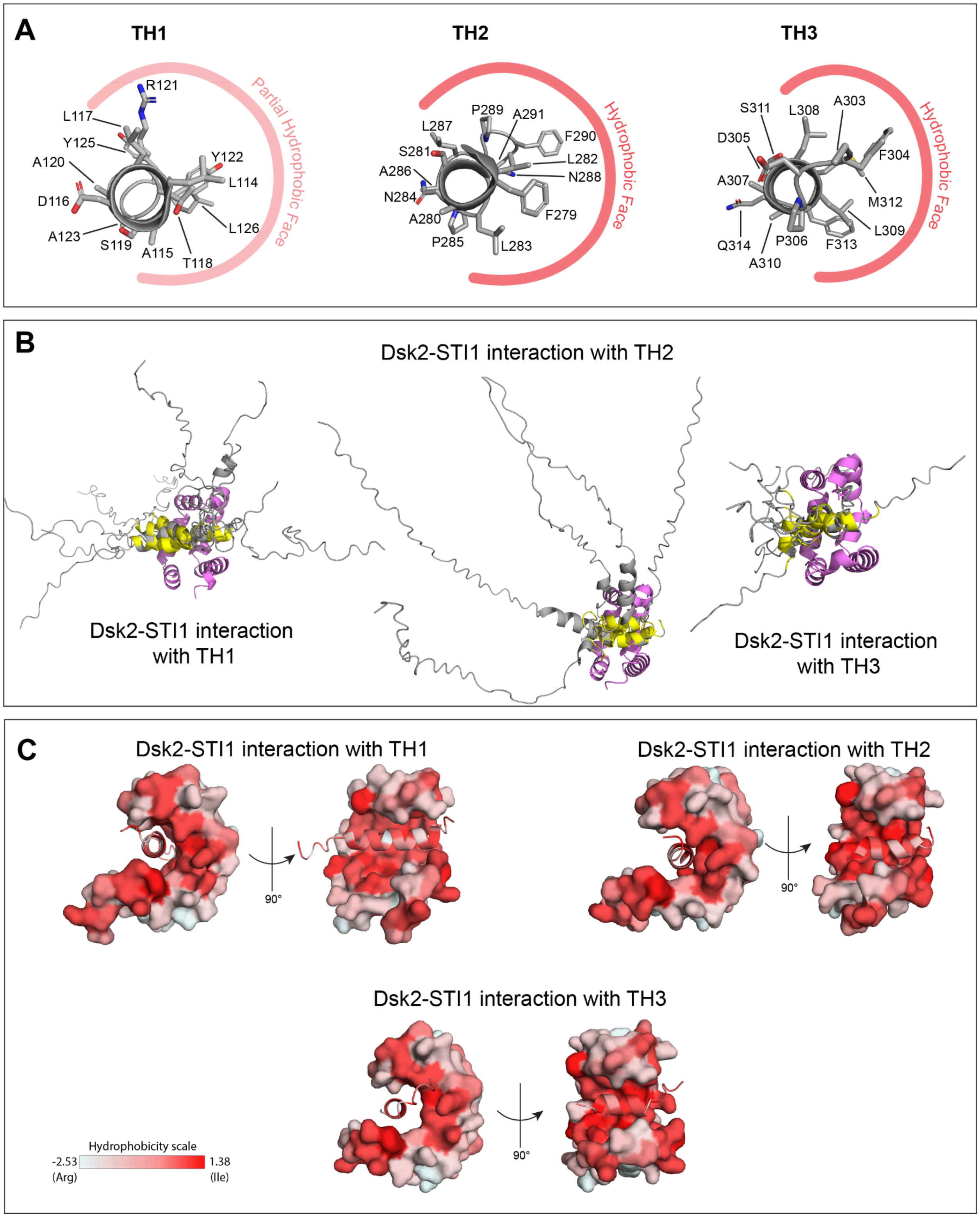

**Figure S10. Hydrophobic interactions between amphipathic transient helices and STI1 domain of Dsk2.** (A) Cartoon representations of TH1, TH2, and TH3 are shown, highlighting the distribution of amino acid residues (shown as sticks, with red, blue, and orange colors representing O, N, and S atoms, respectively) within these transient helices. The TH2 and TH3 helices of Dsk2 are amphipathic, while the TH1 helix has a partial hydrophobic face. Representations are based on AlphaFold-predicted structure of Dsk2 (AF-P48510-F1-v4). (B) Overlays of all five predicted models of AlphaFold2 multimer runs are shown for STI1-TH1, STI1-TH2, and STI1-TH3 (STI1 domain residues 147-228, TH1 region: residues 77-146, TH2 region: residues 229-291, TH3 region: residues 292-325). Residues with CSP > 0.04 ppm between Dsk2 FL and ΔSTI1 (from Figure 4B) are highlighted yellow. (C) highlight the hydrophobic interactions between the STI1 domain (surface representation) and transient helices TH1, TH2, and TH3 (cartoon representation) in Dsk2 using predicted AlphaFold models (alternative representations of the same structures presented in Figure 6). All amino acid residues are colored from white to red based on their increasing hydrophobicity using Pymol.

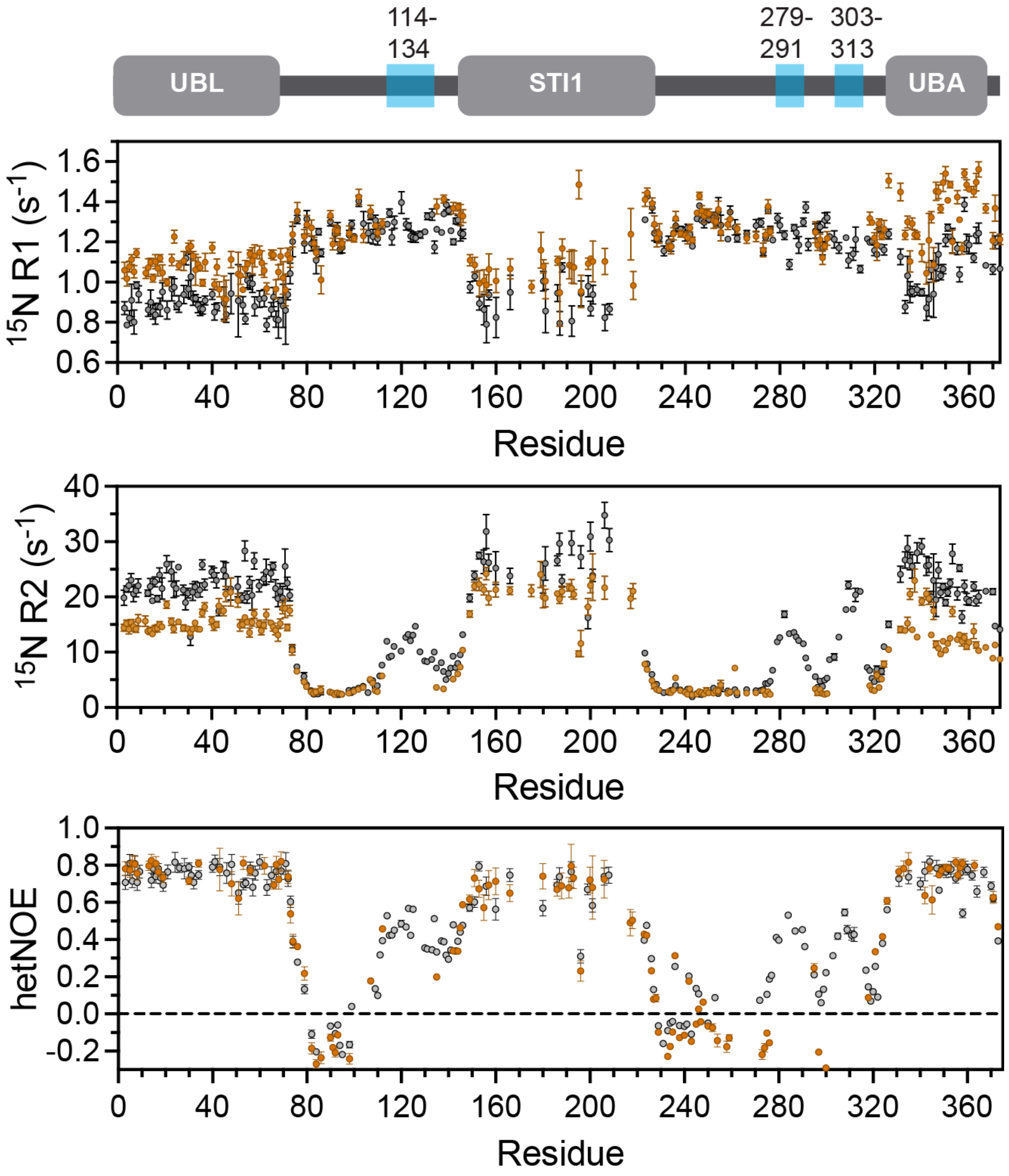

**Figure S11. Comparison of NMR relaxation properties for Dsk2 Δ3TH and Dsk2 FL.** Comparison of $^{15}$N $R_1$ and $R_2$ relaxation rates, and hetNOE values between Dsk2 FL (gray) and Dsk2 Δ3TH (brown). Errors in R1 and R2 relaxation rates were determined using 500 Monte Carlo trials using RELAXFIT. Errors in hetNOE measurements were determined using the SE propagation formula. There is a substantial increase in R1 relaxation rates and corresponding decrease in R2 relaxation rates of Dsk2 Δ3TH in all folded domains (UBL, STI1, and UBA), indicating loosening of structural compactness (has become more flexible) compared to the Dsk2 FL. A comparison of the hetNOE values reveals that deletion of all three transient helical regions has no significant effect on the fast local dynamics in the protein, including within the STI1 domain.

**A**

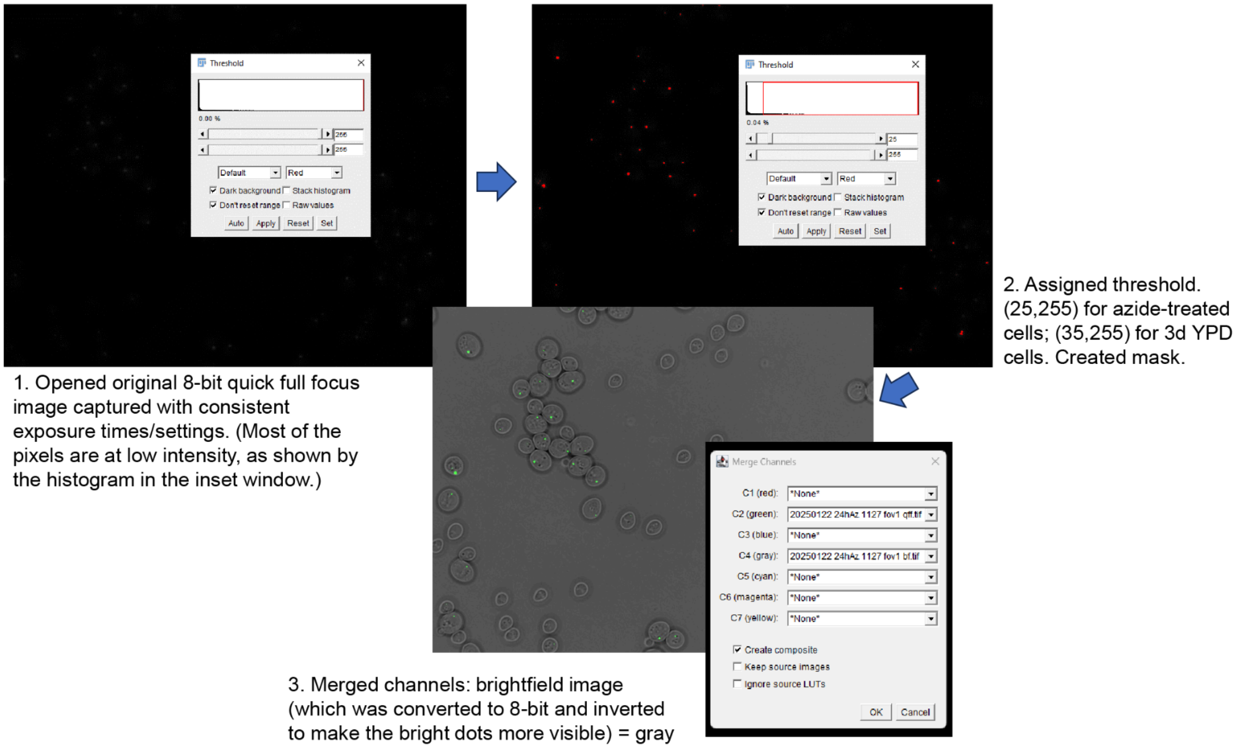

**B**

```
//for  brightfield images
setMinAndMax(0, 65535);
run("8-bit");
run("Invert");

//for quick-full-focus GFP images
run("8-bit")
setThreshold(35, 255, "raw");
run("Convert to Mask");

//for merging the mask and brightfield image for counting
run("Merge Channels..."); //NOTE: make C4 gray the bf image and choose a color for the GFP image (green is easy to see)
run("Stack to RGB");
//then count with Cell Counter tool
```

**Figure S12. Image processing workflow for intensity thresholding.** (A) Example screenshots of the image processing workflow for intensity threshold analysis accompanying Figure 6. (B) Three separate ImageJ macro scripts (file type *.ijm) were used to facilitate the workflow.

**Table S1. Helix parameters for the three transient helices of Dsk2.** All hydrophobic amino acids (in the order of increasing hydrophobicity: A, M, L, and F) have been highlighted in red in each sequence, showing their number and distribution in respective helices.

| Transient helix | Helix length | Sequence | Number of hydrophobic amino acids |
|---|---|---|---|
| TH1 (114-126) | 13 | **LA**D**L**TS**A**RY**A**GY**L** | 6 |
| TH2 (279-291) | 13 | **FA**S**LL**NP**AL**NP**FA** | 8 |
| TH3 (303-314) | 12 | **AF**DP**ALLA**S**MF**Q | 8 |

**Table S2. Amino acid sequence of purified Dsk2 constructs.** Color coding: UBL (blue), transient helical regions (orange), STI1 domain (red), and UBA (violet).

| Construct | Amino acid sequence |
|---|---|
| Dsk2 FL | MSLNIHIKSGQDKWEVNVAPESTVLQFKEAINKANGIPVANQRLIYSGKILKDDQTVESYHIQDGHSVHLVKSQPKPQTASAAGANNATATGAAAGTGATPNMSSGQSAGFNPLADLTSARYAGYLNMPSADMFGPDGGALNNDSNNQDELLRMMENPIFQSQMNEMLSNPQMLDFMIQSNPQLQAMGPQARQMLQSPMFRQMLTNPDMIRQSMQFARMMDPNAGMGSAGGAASAFPAPGGDAPEEGSNTNTTSSSNTGNNAGTNAGTNAGANTAANPFASLLNPALNPFANAGNAASTGMPAFDPALLASMFQPPVQASQAEDTRPPEERYEHQLRQLNDMGFFDFDRNVAALRRSGGSVQGALDSLLNGDV |
| Dsk2 ΔUBL | KPQTASAAGANNATATGAAAGTGATPNMSSGQSAGFNPLADLTSARYAGYLNMPSADMFGPDGGALNNDSNNQDELLRMMENPIFQSQMNEMLSNPQMLDFMIQSNPQLQAMGPQARQMLQSPMFRQMLTNPDMIRQSMQFARMMDPNAGMGSAGGAASAFPAPGGDAPEEGSNTNTTSSSNTGNNAGTNAGTNAGANTAANPFASLLNPALNPFANAGNAASTGMPAFDPALLASMFQPPVQASQAEDTRPPEERYEHQLRQLNDMGFFDFDRNVAALRRSGGSVQGALDSLLNGDV |
| Dsk2 mutUBA | MSLNIHIKSGQDKWEVNVAPESTVLQFKEAINKANGIPVANQRLIYSGKILKDDQTVESYHIQDGHSVHLVKSQPKPQTASAAGANNATATGAAAGTGATPNMSSGQSAGFNPLADLTSARYAGYLNMPSADMFGPDGGALNNDSNNQDELLRMMENPIFQSQMNEMLSNPQMLDFMIQSNPQLQAMGPQARQMLQSPMFRQMLTNPDMIRQSMQFARMMDPNAGMGSAGGAASAFPAPGGDAPEEGSNTNTTSSSNTGNNAGTNAGTNAGANTAANPFASLLNPALNPFANAGNAASTGMPAFDPALLASMFQPPVQASQAEDTRPPEERYEHQLRQLNDMAAFDFDRNVAALRRSGGSVQGALDSLLNGDV |
| Dsk2 Δ3TH | MSLNIHIKSGQDKWEVNVAPESTVLQFKEAINKANGIPVANQRLIYSGKILKDDQTVESYHIQDGHSVHLVKSQPKPQTASAAGANNATATGAAAGTGATPNMSSGQSAGFNPGPDGGALNNDSNNQDELLRMMENPIFQSQMNEMLSNPQMLDFMIQSNPQLQAMGPQARQMLQSPMFRQMLTNPDMIRQSMQFARMMDPNAGMGSAGGAASAFPAPGGDAPEEGSNTNTTSSSNTGNNAGTNAGTNAGANTAANPNAGNAASTGPPVQASQAEDTRPPEERYEHQLRQLNDMGFFDFDRNVAALRRSGGSVQGALDSLLNGDV |
| Dsk2 ΔSTI1 | MSLNIHIKSGQDKWEVNVAPESTVLQFKEAINKANGIPVANQRLIYSGKILKDDQTVESYHIQDGHSVHLVKSQPKPQTASAAGANNATATGAAAGTGATPNMSSGQSAGFNPLADLTSARYAGYLNMPSADMFGPDGGALNNDAGMGSAGGAASAFPAPGGDAPEEGSNTNTTSSSNTGNNAGTNAGTNAGANTAANPFASLLNPALNPFANAGNAASTGMPAFDPALLASMFQPPVQASQAEDTRPPEERYEHQLRQLNDMGFFDFDRNVAALRRSGGSVQGALDSLLNGDV |

| | |
|---|---|
| Dsk2 IDR+STI1 | KPQTASAAGANNATATGAAAGTGATPNMSSGQSAGFNPLADLTSARYAGYLNMPSADMFGPDGGALNNDSNNQDELLRMMENPIFQSQMNEMLSNPQMLDFMIQSNPQLQAMGPQARQMLQSPMFRQMLTNPDMIRQSMQFARMMDPN |
| Dsk2 STI1+IDR | SNNQDELLRMMENPIFQSQMNEMLSNPQMLDFMIQSNPQLQAMGPQARQMLQSPMFRQMLTNPDMIRQSMQFARMMDPNAGMGSAGGAASAFPAPGGDAPEEGSNTNTTSSSNTGNNAGTNAGTNAGANTAANPFASLLNPALNPFANAGNAASTGMPAFDPALLASMFQPPVQASQAEDTR |
| Dsk2 UBL | MSLNIHIKSGQDKWEVNVAPESTVLQFKEAINKANGIPVANQRLIYSGKILKDDQTVESYHIQDGHSVHLVKSQP |
| Dsk2 UBA | RPPEERYEHQLRQLNDMGFFDFDRNVAALRRSGGSVQGALDSLLNGDV |

**Table S3. Molecular weights and molar extinction coefficients of purified Dsk2 constructs used for concentration determination.**

| Construct | Molecular weight (Da) | Molar extinction coefficient ($M^{-1}cm^{-1}$) |
|---|---|---|
| Dsk2 FL | 39345 | 12950 |
| Dsk2 ΔUBL | 30969 | 4470 |
| Dsk2 mutUBA | 39283 | 12950 |
| Dsk2 Δ3TH | 34178 | 9970 |
| Dsk2 ΔSTI1 | 30007 | 12950 |
| Dsk2 IDR+STI1 | 15945 | 2980 |
| Dsk2 STI1+IDR | 19106 | NA* |

*NA: Not applicable*
**Concentration of Dsk2 STI1+IDR was estimated by SDS-PAGE gel using concentration standards of similar molecular weight protein.*

**Table S4. Yeast strain list**

| Strain | Genes manipulated | Simplified name | Figures | Ref |
|---|---|---|---|---|
| sJR1255 | *rpn1*::*RPN1-GFP* (HIS3) | WT | 6, S7, S8 | a |
| sJR1127 | *rpn1*::*RPN1-GFP* (HIS3) *rad23*Δ*::*KanMX | *rad23*Δ | 6, S7, S8 | a |
| sJR2659 | *rpn1*::*RPN1-GFP* (HIS3) DSK2 Δ145–223 | *DSK2* Δ*STI1* | 6, S7, S8 | b |
| sJR2660 | *rpn1*::*RPN1-GFP* (HIS3) *rad23*Δ*::* KanMX DSK2 Δ145–223 | *rad23*Δ *DSK2* Δ*STI1* | 6, S7, S8 | b |

Background of all strains: MATα his3Δ1 leu2Δ0 lys2Δ0 ura3Δ0 (BY4742)

a. (Waite *et al*, 2024)
b. This study

**Table S5. Plasmid and repair DNA used for generating Dsk2 ΔSTI1 yeast strain**

| **Plasmid** | | |
|---|---|---|
| pJR1173 | Extrachromosomal plasmid expressing *S. pyogenes* cas9, sgRNA used to delete the STI1 domain of Dsk2, and yeast *LEU2*. Derived from Addgene plasmid #67639 (pML107, (Laughery *et al*, 2015)) | Guide RNA contained the yeast Dsk2 targeting sequence: 5'-TGTAGCATTTGCCTGGCTTG-3', which is followed in the yeast genome by a PAM sequence 'TGG' at the 3' end. |
| **Repair DNA** | | |
| pRL1528 | Repair duplex DNA for deletion of the Dsk2 STI1 domain via homologous recombination (only top strand shown) | CAATCCGCTGGCCGACTTGACCAGTGCCAGATACGCTGGATATTTGAATATGCCATCTGCAGACATGTTTGGCCCGGACGGTGGTGCATTAAACAACGACGCCGGTATGGGCTCTGCAGGTGGGGCTGCCTCTGCCTTCCCCGCTCCTGGTGGCGATGCTCCAGAGGAAGGCTCCAACACGAACACTACTTCCTCATCCAACACAGGGAACAACGCAGG |

**Table S6. SAXS data collection**

| | |
|---|---|
| *(a)* Sample details | |
| Organism | *S. cerevisiae* |
| Source (Catalogue No. or reference) | Expressed in *E. coli* (this work) |
| Description: sequence (including Uniprot ID + uncleaved tags), bound ligands/modifications, etc. | Full-length Dsk2 (no tags), Uniprot ID P48510 |
| Extinction coefficient ε in $M^{-1}$ $cm^{-1}$ (wavelength in nm) | 12950 (280) |
| Molecular mass M from chemical composition (Da) | 39,300 |
| For SEC-SAS, loading volume/concentration (mg $ml^{-1}$), injection volume (μl), flow rate (ml $min^{-1}$) | 6.11, 300, 0.65 |
| Solvent composition and source | pH 6.8 20 mM NaPhos, 0.5 mM EDTA, 0.02% $NaN_3$ |
| *(b)* SAS data collection parameters | |
| Instrument | SIBYLS facility (beamline 12.3.1) at the Advanced Light Source, with Pilatus X3 2M detector (Dectris) |
| Wavelength (Å) | 1.240 |
| Camera length (m) | 2.077 |
| *q*-measurement range | 0.0114-0.0473 |
| Normalization | Transmitted intensity |
| Exposure time/number | 2.0 seconds (600 frames) |

| | |
|---|---|
| Sample Configuration | SEC-MALS-SAXS using a Shodex KW-803 and an Agilent 1260 Series HPLC. UV data was measured with an Agilent 1290 DAD, and MALS/RI data by DAWN HELEOS-II (18-angle) and Optilab T-rEX (RI) instruments (Wyatt Technology). SAXS data was measured in a 1 mm, Mica-windowed flow cell |
| Sample Temperature | 25 °C |
| *(c)* Software employed | |
| SAXS data reduction | Radial averaging; frame comparison, averaging, and subtraction done using BioXTAS RAW 2.1.1 (Hopkins *et al*, 2017) |
| Basic analysis: Guinier, M.W. P(r) | Guinier fit and M.W. using BioXTAS RAW. RAW uses MoW and Vc M.W. methods (Rambo & Tainer, 2013; Piiadov *et al*, 2019) |
| MALS-RI analysis | Astra 7 (Wyatt) |
| *(d)* Structural Parameters | |
| Guinier Analysis | Full-length Dsk2 |
| I(0) | 62.05 ± 0.25 |
| $R_g$ (Å) | 35.13 ± 0.18 |
| q-range ($Å^{-1}$) | 0.01444 – 0.03725 |
| Quality-of-fit parameter (with definition) | 0.9952 ($r^2$) |
| *M* from MALS (kDa) | 35 |

**Supplementary Movies**

**Movie S1:** Representative CALVADOS molecular dynamics simulation for full-length Dsk2 where UBL and UBA domains are left unrestrained. Colors are blue (UBL), red (STI1), purple (UBA), orange (segments corresponding to TH1, TH2, and TH3 regions). The movie frame rate corresponds to 100 ps per second.

**Movie S2:** Representative CALVADOS molecular dynamics simulation for full-length Dsk2 where UBL and UBA domains are restrained in the bound conformation (see Methods). Colors are the same as described in Movie S1. The movie frame rate corresponds to 100 ps per second.
